# Lipid droplet accumulation drives glutamine-dependent NLRP3 inflammasome activation

**DOI:** 10.64898/2026.01.13.699337

**Authors:** Najd M. Aljadeed, Erato Pambori, Yujia Zhang, Greg Austin, Antoni Olona, Guido Franzoso, Shamith Samarajiwa, Alejandra Tomas, Frederick Wai Keung Tam, Paras K. Anand

## Abstract

The NLRP3 inflammasome is implicated in the pathogenesis of many inflammatory and metabolic diseases. Although metabolic cues can shape NLRP3 inflammasome activation, the role of lipid metabolism in this response is poorly understood. Lipids are essential for energy and redox homeostasis, membrane biogenesis, and diverse cellular processes. However, excessive accumulation of lipids such as free fatty acids can be highly cytotoxic. Consequently, surplus free fatty acids are esterified into triglycerides and cholesteryl esters, which are then stored within lipid droplets (LDs). Interestingly, transcriptomic data analyses revealed that macrophages exposed to inflammasome-activating stimulus undergo a broad metabolic rewiring favouring lipid storage, suggesting a link between LDs and inflammasome response. Here, we show that LD accumulation actively promotes NLRP3 inflammasome activation. Increasing LD abundance, either by fatty acid supplementation or by inhibition of lipolysis, led to robust caspase-1 activation and IL-1β release both *in vitro* and *in vivo*. Mechanistically, LD expansion promoted cellular reorganisation resulting in the formation of peri-droplet mitochondria, a distinct mitochondrial subpopulation which exhibited elevated membrane potential and increased ATP-linked respiration. Furthermore, LD-rich macrophages underwent metabolic reconfiguration whereby they exhibited a pronounced reliance on pyruvate and glutamine metabolism, but not fatty acid oxidation, to sustain mitochondrial bioenergetics. Surprisingly, inflammasome activation was found dependent predominantly on glutaminolysis, highlighting a key role for glutamine catabolism in LD-rich cells. Collectively, these findings identify a glutamine-dependent pathway as a key driver linking excess lipid storage to NLRP3 inflammasome, with profound implications in metabolic diseases.

## Introduction

Metabolic diseases such as type-2 diabetes and obesity are among the most significant growing threats to global public health. The causes of these diseases are multi-factorial and generally involve a combination of lifestyle factors, genetic predispositions, and environmental influences. Irrespective of the heterogeneous and complex aetiology of metabolic diseases, it is widely recognised that a chronic state of low-grade inflammation, often referred to as meta-inflammation, is a common driver of metabolic disease progression, underscoring the intimate interplay between nutritional imbalance and immune dysfunction^1^. A central mediator of metabolic inflammation is the NLRP3 inflammasome, a cytoplasmic multiprotein complex that senses metabolic danger signals and triggers the release of pro-inflammatory cytokines IL-1β and IL-18, as well as the induction of inflammatory cell death, pyroptosis^2–5^. This has prompted growing interest in understanding how metabolic signals regulate inflammation in disease.

Lipid metabolism plays a crucial role in immune cell function, requiring a precise regulation of lipid biosynthesis, storage, and breakdown depending on cellular demands ^6–8^. Fatty acids can be generated *de novo* or imported from extracellular sources before being incorporated into triglycerides, phospholipids, and other complex lipids in a process known as lipogenesis^9–11^. This route enables excess free fatty acids to be stored in lipid droplets (LDs), which sequester neutral lipids within a phospholipid monolayer. During nutrient scarcity, stored triglycerides are hydrolysed in a process known as lipolysis, primarily mediated by adipose triglyceride lipase (ATGL)^12^. ATGL activity liberates fatty acids from LDs, which are subsequently transported to mitochondria for β-oxidation^13^. Dysregulation of any of these pathways perturbs lipid homeostasis and has been implicated in the pathogenesis of both metabolic and chronic inflammatory disorders^14–16^.

Although once considered as inert lipid reservoirs that merely sequester free fatty acids to prevent lipotoxicity, LDs are now recognised as dynamic organelles with important regulatory functions in immunity and metabolism^17^. Emerging evidence indicates that LDs can modulate inflammatory signalling depending on the extent of lipid mobilisation, their composition of lipid species, and their interactions with cellular organelles^18^. For example, inhibition of triglyceride synthesis, which limits LD formation, has been shown to suppress prostaglandin synthesis and cytokine secretion in macrophages^19^. Conversely, the inhibition of lipolysis, which promotes LD abundance, can similarly attenuate prostaglandin production and IL-6 secretion^20^. Although the precise mechanisms underpinning these activities remain largely unknown, the above observations substantiate LDs as active regulators, rather than passive bystanders of immune responses.

Notably, previous studies have shown that the aberrant accumulation of LDs in immune cells is a hallmark of numerous metabolic diseases ^21,22^. However, this phenomenon has often been dismissed as a downstream consequence of the disease state, rather than a causative contributor to disease pathogenesis. In support of latter hypothesis, in microglia, LD abundance has been shown to drive a dysfunctional pro-inflammatory state that accompanies neurodegeneration^23^, while LD-rich immune cells have been shown to exert pathogenic functions in atherosclerosis and allergic inflammation^24,25^. Collectively, these results support the notion that LD accumulation can shape immune responses in a context-dependent manner influenced by both metabolic and inflammatory cues.

While LDs are most abundant in adipocytes, they are present in nearly all cell types, where they interact dynamically with other organelles to regulate lipid trafficking and energy and redox metabolism^26,27^. Among cellular organelles, mitochondria have emerged as a key compartment associated with LDs, giving rise to a distinct subpopulation known as peri-droplet mitochondria (PDMs). PDMs exhibit bioenergetic properties that are distinct from those of other cytoplasmic mitochondria. In particular, LD-mitochondria interactions have been implicated in cellular responses to inflammatory stimuli^27^. Interestingly, the NLRP3 inflammasome is especially sensitive to perturbations in mitochondrial function^28,29^; yet, whether LDs, or their interaction with mitochondria, directly regulate NLRP3 activation remains unexplored.

Here, we demonstrate that LD accumulation potentiates NLRP3 inflammasome activation. We demonstrate that enhanced LD abundance, either through oleic acid (OA) supplementation or by inhibition of lipolysis, results in robust caspase-1 activation, both *in vitro* and *in vivo*. Mechanistically, LD expansion accompanied increased formation of PDMs, elevated mitochondrial membrane potential (ΔΨm), and enhanced ATP-linked respiration in metabolic flux assays. We find that LD-rich cells display a shift in substrate preference, exhibiting a marked dependence on pyruvate and glutamine oxidation to sustain mitochondrial bioenergetics. Remarkably, fatty acid utilisation remained unchanged. Surprisingly, we identify glutamine oxidation as the key metabolic driver of inflammasome activation in LD-rich cells. Together, these findings demonstrate that LD abundance and PDM formation engage glutamine metabolism to potentiate NLRP3 inflammasome activation, offering novel mechanistic insight into lipid-driven inflammation in metabolic disease.

## Results

### NLRP3 inflammasome-activating stimuli remodel pathways involved in lipid storage

To identify metabolic pathways altered during NLRP3 inflammasome activation, we first performed transcriptomic analysis of bone marrow–derived macrophages (BMDMs) stimulated with NLRP3 inflammasome stimulus, LPS+ATP (see Methods). Gene Ontology (GO) enrichment analysis of all differentially expressed genes revealed that, in addition to immune-related processes such as defence response, cell death, and cytokine response, several additional pathways were significantly enriched **(Fig. 1A)**. Notably, the term “response to lipid” emerged as one of the top enriched categories distinctively in macrophages exposed to LPS+ATP, but not in cells exposed to LPS alone **(Fig. 1A, S1)**. This suggested a strong association between lipid metabolism and NLRP3 inflammasome activation.

**Figure 1.**
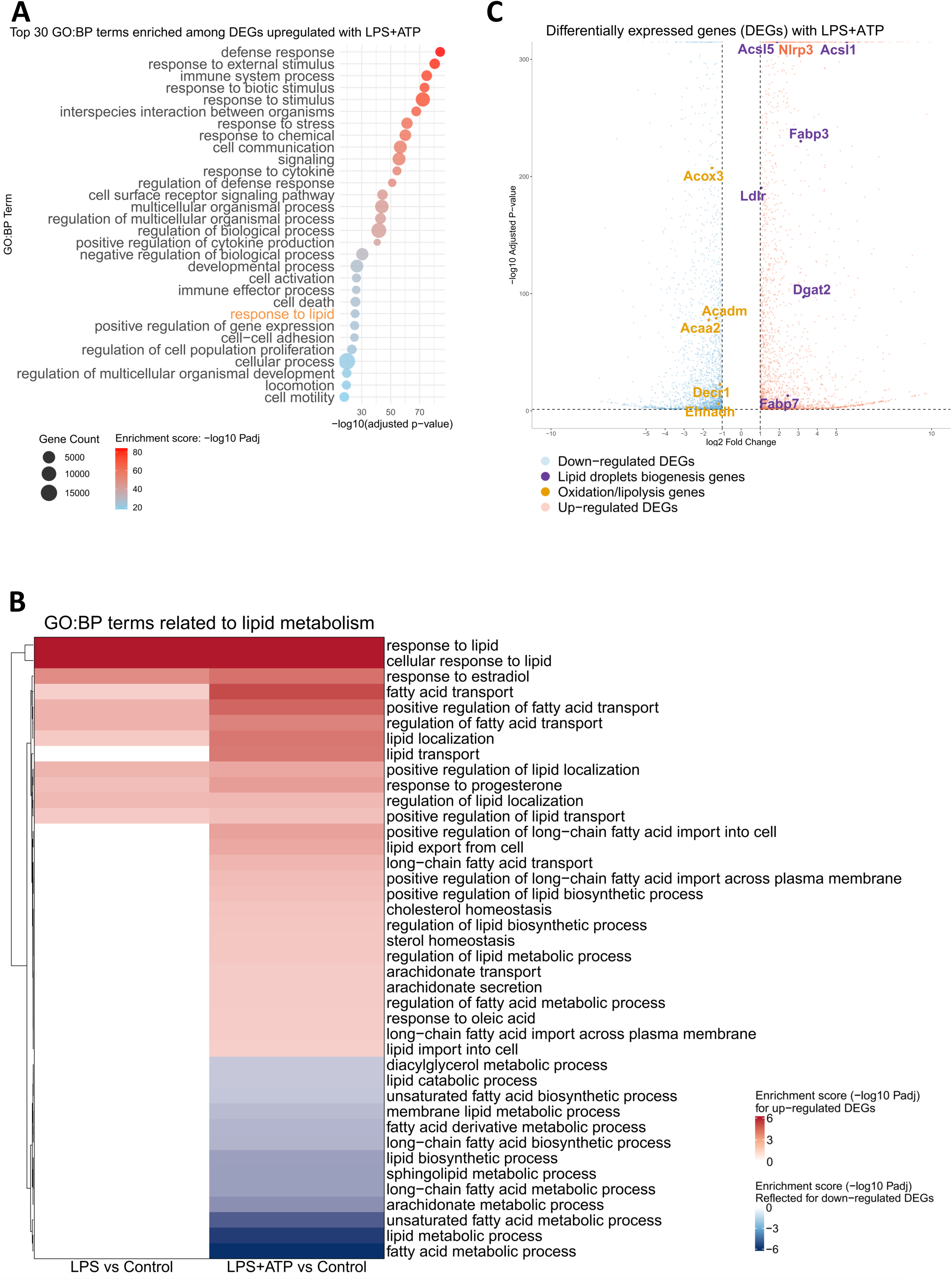
NLRP3 inflammasome-activating stimuli remodel pathways involved in lipid storage. **(A)** Top 30 enriched Gene Ontology Biological Process (GO:BP) terms among upregulated DEGs in LPS+ATP treated versus control BMDMs, identified by g:Profiler and summarized with rrvgo. Enrichment score = -log_10_ (FDR-adjusted p value); gene count = number of DEGs within each GO:BP term. Full results are provided in Supplementary Materials. **(B)** Heatmap of lipid metabolism-related GO:BP terms enriched in DEGs from LPS versus control and LPS+ATP versus control macrophages. Terms enriched in upregulated DEGs are shown in red; downregulated terms are assigned negative enrichment scores and shown in blue. **(C)** DEGs from LPS+ATP versus control and LPS versus control macrophages, with genes related to lipid droplet biogenesis highlighted in purple and genes related to lipid oxidation and lipolysis highlighted in orange.

To further delineate this relationship and narrow down on the key pathways that may regulate the NLRP3 inflammasome, we selectively examined GO pathways associated with lipid response **(Fig. 1B)**. This analysis revealed coordinated transcriptional reprogramming leading to the upregulation of genes involved in lipid biosynthesis, import, and localization accompanied by the downregulation of genes related to lipid catabolism **(Fig. 1B)**. Overall, macrophages exposed to inflammasome stimulation displayed a distinct lipid-related gene expression signature favouring lipid storage when compared to LPS alone.

Since the observed transcriptional changes in lipid metabolism pathways could reflect upstream regulatory events that potentiate inflammasome activation, we next examined the expression of individual genes within these pathways **(Supplementary Files 1-7).** In line with the lipid remodelling observed above, there was a coordinated induction of genes associated with lipid synthesis and accumulation, which suggested that lipid droplet (LD) formation may play an active role in inflammasome activation. Notably, several genes involved in LD biogenesis and fatty acid esterification including *Dgat2*, *Acsl1*, and *Fabp3*^30,31^ were significantly upregulated. Conversely, a subset of genes implicated in lipid oxidation and lipolysis, including *Acox3, Acadm, and Acaa2* ^32,33^ were downregulated, suggesting a metabolic shift towards lipid retention and enhanced LD stability under inflammasome-activating conditions **(Fig. 1C)**. Together, these transcriptional changes implicate LD abundance as an active process that contributes to NLRP3 inflammasome activation prompting us to investigate this link further.

### LD accumulation results in increased NLRP3 inflammasome activation and pyroptosis

LDs are highly dynamic organelles composed of a neutral lipid core enclosed by a phospholipid monolayer^6^. While adipocytes are uniquely adapted, non-adipocyte cells such as macrophages can also accumulate LDs under conditions that favour lipid excess. Recent studies have suggested that LDs may modulate inflammatory signalling^19,20^. However, whether LDs directly modulate inflammasome activation and the mechanism involved has not been explored. Given that our transcriptomic findings indicated enhanced expression of genes promoting LD biogenesis when exposed to NLRP3 stimulus, we hypothesised that LDs could actively contribute to inflammasome activation.

To probe this, we first determined the basal levels of LDs in mouse BMDMs by labelling cells with BODIPY (493/503), which upon binding to neutral lipids emits a green fluorescence signal^34^. Control untreated cells labelled with BODIPY showed few LDs **(Fig. 2A**, *left panels*). We next supplemented cells with monounsaturated oleic acid (OA; C18:1), a well-established inducer of LD expansion in various cell types^35^. OA enters lipid biosynthesis pathway undergoing esterification which leads to triglyceride synthesis and storage within LDs^36^. In agreement with previous studies, exposure to OA significantly increased LD numbers within macrophages **(Fig. 2A**, *right panels*). LD abundance expanded with increasing OA concentrations demonstrating the versatility of this model to examine the effect of LD accumulation on inflammasome activation **(Fig. 2B)**.

**Figure 2.**
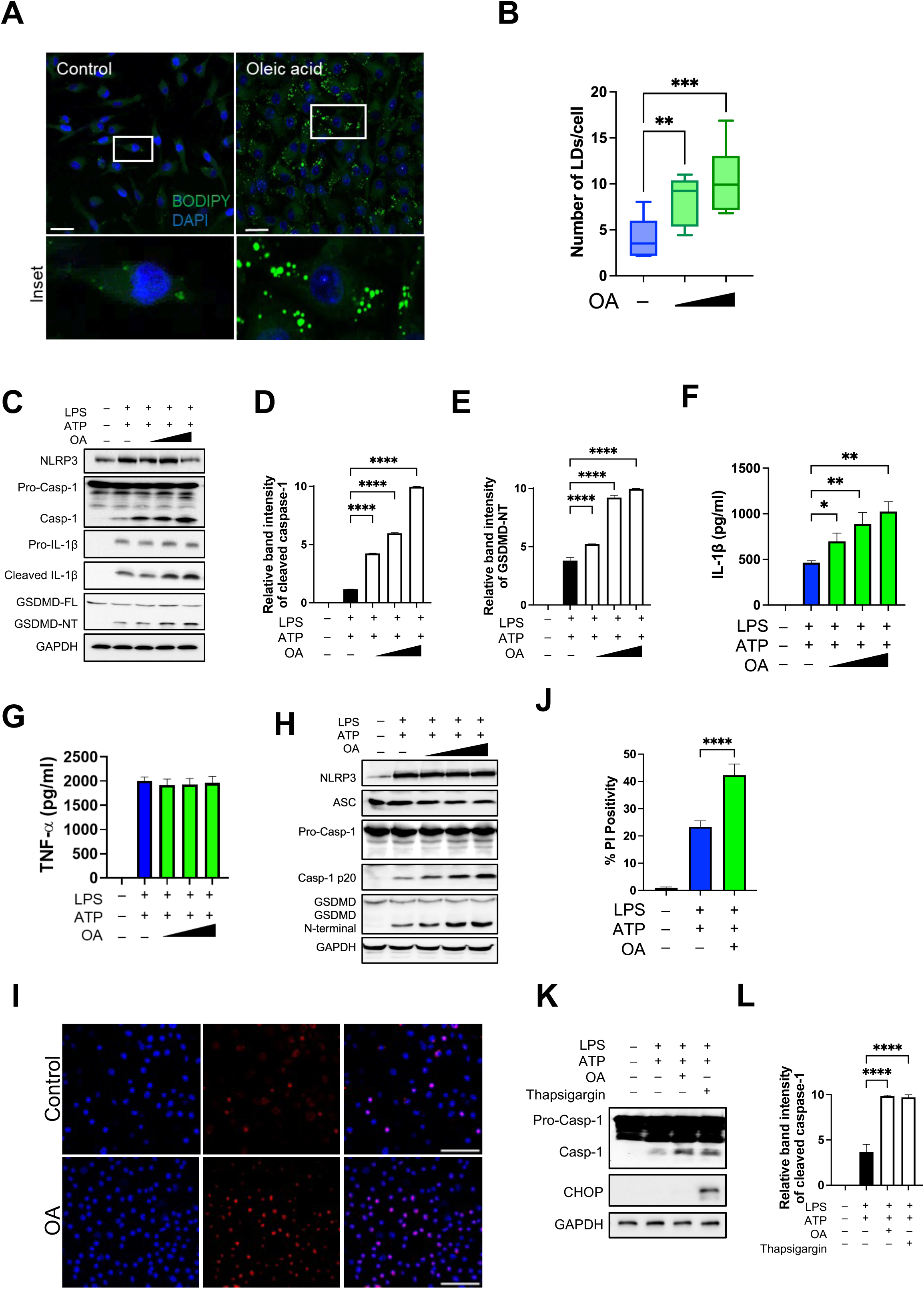
LD accumulation results in increased NLRP3 inflammasome activation and pyroptosis. **(A)** Confocal microscopy images of BMDMs grown in the presence of oleic acid (OA; 90µM) for 24 h followed by incubation with BODIPY 493/503 (2 µM for 20 min in PBS) at 37°C. **(B)** Quantitative analysis of the number of lipid droplets (LDs) per cell at increasing concentrations (60µM, 90µM) of OA. **(C)** BMDMs stimulated with inflammasome priming signal LPS (500 ng/ml; 3.5 hours) were exposed to increasing concentrations of OA (60, 90, and 150 µM) for 24 h prior to exposure to ATP (5 mM; 30 min). Cell lysates were immunoblotted with the indicated antibodies. **(D, E)** Relative band intensity of cleaved caspase-1 as measured by ImageJ. **(F)** IL-1β and TNF-α released from cells treated as above. **(H)** Immortalised BMDMs (iBMDMs) stimulated with inflammasome priming signal LPS (500 ng/ml; 3.5 hours) were exposed to increasing concentrations of OA (30, 60, and 90 µM) for 48 h prior to exposure to ATP (5 mM; 30 min). **(I)** Fluorescent images from iBMDMs grown in the presence of OA and exposed to LPS+ATP followed by propidium iodide staining. **(J)** Percentage of propidium iodide (PI) positive cells from experiment in (I). **(K)** iBMDMs exposed to LPS and ATP in the presence or absence of OA (30 µM; 48 h) or thapsigargin (10 µM; 5 h). Cell lysates were immunoblotted with the indicated antibodies. **(L)** Relative band intensity of cleaved caspase-1 as measured by ImageJ. Data shown are mean ± SD. Experiments shown are representative of at least three independent experiments. Panel A, scale bars, 20 μm, Panel I, scale bars, 50 μm. *, p<0.05; **, p < 0.01; ***, p < 0.001; ****, p < 0.0001, by one-way ANOVA, or Student’s *t* test.

Next, we determined whether elevated LD abundance regulates inflammasome activation. To this effect, primary BMDMs were grown in the presence of OA for 24 hours before exposing them to LPS and ATP. Control macrophages exposed to LPS and ATP demonstrated cleavage of 45 kDa pro-caspase-1 to its 20 kDa active form. However, the processed active form of caspase-1 significantly increased in a dose-dependent manner in OA-fed cells revealing elevated NLRP3 inflammasome activation **(Fig. 2C, D)**. Consequently, compared to control macrophages, the secretion of caspase-1-dependent cytokine IL-1β also exhibited a significant increase while the inflammasome-independent cytokine TNF-α remained unchanged **(Fig. 2F, G, Fig. S2)**. Similar results were obtained with immortalised BMDMs though notably at a much lower OA concentration (30 µM vs 90 µM for primary BMDMs) **(Fig. 2H)**. The expression of sensor protein NLRP3 and the adaptor protein ASC remained unaltered in cells supplemented with OA suggesting that the increase in inflammasome activation is independent of NLRP3 priming **(Fig. 2C, H)**.

Active caspase-1 further cleaves gasdermin D (GSDMD), with the N-terminal of GSDMD translocating to the plasma membrane to form pores and induces an inflammatory cell death, pyroptosis^37,38^. In agreement with elevated caspase-1, LD accumulation resulted in a dose-dependent increase in GSDMD N-terminal fragment as compared to control cells **(Fig. 2C, E, H)**. Furthermore, the percentage of cells positive for propidium iodide (PI) staining, a membrane-impermeable dye, was significantly higher in cells with LDs compared to control cells **(Fig. 2I, J)** suggesting a reduced membrane integrity in these cells.

The increase in inflammasome activity was not limited to the P_2_X7R agonist and inflammasome stimulus, ATP. Exposure to nigericin similarly resulted in increased IL-1β secretion by oleate-loaded cells **(Fig. S3)**. NLRP3 activation can be triggered by both soluble and particulate ligands^39^. The increase in NLRP3 activation in LD-rich macrophages extended to particulate stimuli. Exposure to alum similarly elevated inflammasome activation in LD-accumulating conditions **(Fig. S3)**. These data suggested that NLRP3 activation by LDs is independent of the nature of the upstream NLRP3 stimulus.

Disruption in lipid homeostasis is strongly associated with ER stress, a cellular mechanism which activates the unfolded protein response (UPR) aimed at restoring homeostasis^40^. Some UPR is beneficial but uncontrolled response may trigger apoptosis and inflammatory signalling via the upregulation of the C/EBP homologous protein (CHOP)^40^. Consequently, we next assessed whether OA supplementation triggered UPR response by examining CHOP expression in cell lysates. In parallel, we supplemented cells with thapsigargin, a specific inhibitor of sarco/endoplasmic reticulum Ca2+ -ATPase (SERCA) pump which robustly activates the UPR, and consequently CHOP expression, by depleting calcium stores in the ER. As expected, exposure to OA or thapsigargin resulted in significant caspase-1 activation **(Fig. 2K, L)**. However, CHOP expression was observed only with thapsigargin exposure while supplementation with OA resulted in no detectable CHOP expression by immunoblotting **(Fig. 2K)**.

Remarkably, our results indicate that the induction of caspase-1 is influenced by the type of fatty acid available to cells. While monounsaturated OA promoted caspase-1, supplementing cells with either saturated palmitic acid (C16:0) or the polyunsaturated arachidonic acid (C20:4) resulted in no significant increase in cleaved caspase-1 expression or IL-1β secretion **(Fig. S4A-C)**. Although different fatty acids have been shown to induce LDs in various cell types, the degree of LD abundance seems to vary^41^. To determine whether palmitic acid or arachidonic acid, like OA, induced LD abundance, mouse BMDMs were labelled with BODIPY. Exposure to palmitic acid and arachidonic acid within macrophages resulted in little to no effect in LD number **(Fig S4D, E)**. These data indicate that a certain threshold of LD abundance is required to trigger enhanced NLRP3 inflammasome activation.

We next examined whether LD accumulation broadly regulates inflammasome activation. To assess this, we infected LD-rich cells with *Salmonella typhimurium* to evaluate the impact on NLRC4 inflammasome activation^42^. In contrast to the NLRP3 inflammasome, cells fed increasing concentrations of OA and infected with *Salmonella* exhibited no significant change in caspase-1 activation compared to control cells infected with *Salmonella* without OA. In agreement, oleate-supplemented cells infected with *Salmonella* also demonstrated cleavage of GSDMD, and IL-1β and TNF-α secretion comparable to that of cells infected with *Salmonella* alone **(Fig. S5)**. Activation of the DNA sensing AIM2 inflammasome similarly remained unaltered by LD abundance in response to poly(dA:dT) transfection^43^ **(Fig. S5)**. These data imply that LD accumulation specifically activates the NLRP3 inflammasome.

### Inhibition of lipolysis elevates NLRP3 inflammasome activation in vitro and in vivo

The hydrolysis of stored triglycerides within LDs is dependent on ATGL, a key enzyme in lipolysis^44^. The ATGL enzyme is primarily localised on the surface of LDs where it interacts with regulatory proteins to control lipid mobilisation. Accordingly, inhibition of ATGL blocks lipolysis resulting in cells with elevated LDs. In order to further validate our results, we took an alternative approach of LD expansion by blocking LD turnover with an ATGL inhibitor. In agreement with our earlier data, exposure to the ATGL inhibitor resulted in increased LD abundance compared to control macrophages **(Fig. 3A, S6A)**. Moreover, in agreement with our results, ATGL inhibitor resulted in increased inflammasome activation as shown by increased levels of cleaved caspase-1 by immunoblotting **(Fig. 3B)**. Furthermore, caspase-1 cleavage accompanied increased expression of GSDMD N-terminal fragment and elevated levels of secreted IL-1β **(Fig. 3B, C)** while the levels of TNF-α remained unchanged in the presence of the inhibitor **(Fig. 3D)**. These results indicate that LD accumulation is intricately tied to elevated NLRP3 inflammasome activation.

**Figure 3.**
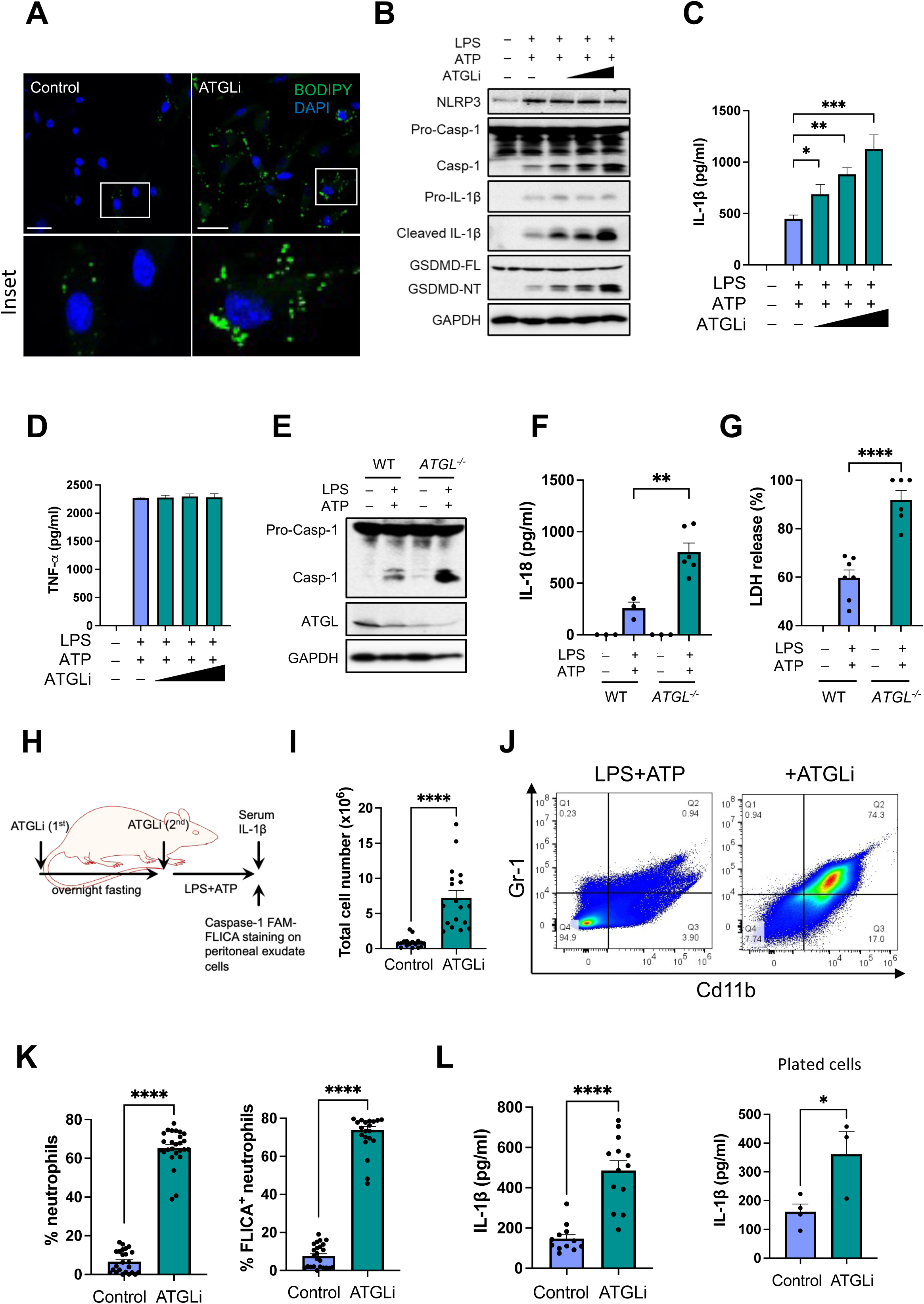
Inhibition of lipolysis elevates NLRP3 inflammasome activation in vitro and in vivo. **(A)** Confocal microscopy images of BMDMs grown in the presence or absence of ATGL inhibitor (ATGLi; 40 µM; 24 h) followed by incubation with BODIPY 493/503 (2 µM for 20 min in PBS) at 37°C. **(B)** BMDMs stimulated with inflammasome priming signal LPS (500 ng/ml; 3.5 hours) were exposed to increasing concentrations of ATGLi (40, 80, and 120 µM) for 24 h prior to exposure to ATP (5 mM; 30 min). Cell lysates were immunoblotted with the indicated antibodies. **(C, D)** IL-1β and TNF-α released from cells treated as above. **(E)** PMA-differentiated WT and ATGL^−/−^ THP-1 macrophages were exposed to LPS (500 ng/ml; 3.5 h) and ATP (5 mM; 30 min). Cell lysates were immunoblotted with the indicated antibodies. **(F, G)** IL-18 and LDH released from cells treated as above. **(H)** Schematic for the *in vivo* experiment: C57BL/6 mice were administered intraperitoneally (i.p.) either with vehicle control (n=12) or ATGLi (50 mg kg−1; n=13) and fasted for 16 h. Next day, mice were again administered either vehicle control or ATGLi and LPS (100 μg kg−1). After 4 h, mice were i.p. injected with ATP for 25 min. **(I)** Quantitative analysis of total peritoneal exudate cell numbers. **(J)** Flow cytometry plots showing gating strategy for CD11b+Gr1+ neutrophils and CD11b+Gr1-monocytes in the above population. **(K)** Frequency of total CD11b+Gr1+ neutrophils *(left panel)* and caspase-1 active neutrophils *(right panel)* in peritoneal exudate cells. **(L)** IL-1β levels in peritoneal exudate *(left panel)* and in the supernatant *(right panel)* 24 h after culturing total peritoneal exudate cells after 24 h. Data shown are mean ± SD. Experiments shown are representative of at least three independent experiments. Panel A, scale bars, 20 μm. *, p<0.05; **, p < 0.01; ***, p < 0.001; ****, p < 0.0001, by one-way ANOVA, or Student’s *t* test.

To corroborate our findings using genetic tools, we generated human THP1 monocytes lacking ATGL by CRISPR/Cas9 gene editing. Analysis of ATGL protein expression on the pooled population of edited cells revealed greater than 90% reduction in ATGL expression compared to control PMA-differentiated WT THP-1 macrophages **(Fig. 3E, S6C)**. When exposed to the NLRP3 stimulus ATP, *ATGL^−/−^* THP-1 macrophages revealed significantly increased caspase-1 activation compared to control THP-1 cells **(Fig. 3E)**. This accompanied increased IL-18 secretion and LDH release by *ATGL^−/−^* cells compared to WT THP-1 macrophages **(Fig. 3F, G)**.

To examine the physiological role of LD expansion during inflammasome activation, we next used a murine model of acute peritonitis^45^. C57BL6/J mice were intraperitoneally injected with the ATGL inhibitor two times followed by LPS. Four hours post-LPS administration, the NLRP3 inflammasome was activated by the intraperitoneal administration of ATP. Peritoneal exudate cells and serum were harvested 25 minutes after ATP administration **(Fig. 3H)**. In animals administered ATGL inhibitor, the total number of cells recruited to the peritoneal cavity was significantly increased compared to animals only given LPS and ATP without lipolysis inhibition **(Fig. 3I)**.

In acute peritonitis, neutrophil recruitment to the peritoneal cavity is dependent on the NLRP3 inflammasome^46,47^. These neutrophils secrete higher levels of IL-1β in peritoneal lavage fluid which is dampened in *Nlrp3*-deficient animals^46^. We next labelled the harvested peritoneal cells with anti-CD11b and anti-Gr1 antibodies to identify and characterize different immune cell populations recruited to the peritoneum. In animals treated with the ATGL inhibitor, there was a marked increase in CD11b+Gr1+ neutrophils within the peritoneal cavity compared to control animals **(Fig. 3J, K)**. Consistent with our study, a higher frequency of caspase-1 positive neutrophils, as measured by FAM-FLICA labelling, were observed following ATGL inhibition **(Fig. 3K**, *right panel*). An increased number of CD11b+Gr1-monocytes were also observed with ATGL inhibitor **(Fig. S6D)**. However, consistent with a role for neutrophils in this model, an increase in caspase-1 positive monocytes was not observed **(Fig. S6E).** Moreover, increased IL-1β was detected in the peritoneal exudate, and peritoneal exudate cells continued to release higher levels of IL-1β for up to 24 hours **(Fig. 3L)**. These data further corroborate a function for LDs in inflammasome activation *in vitro* and *in vivo*.

### LD accumulation promotes increased peri-droplet mitochondria

Under expanded LDs, cells undergo substantial metabolic reconfiguration which primarily depends on establishing close physical interactions between LDs and mitochondria. These contacts give rise to a subset of mitochondrial population, known as peri-droplet mitochondria (PDMs), which form at the lipid droplet interface and are distinct from the broader cytosolic mitochondrial network^48^. Beyond their role in supporting LD expansion, PDMs have been suggested to regulate immune responses^49^. As such, we first explored the functional link between LDs and mitochondria in conditions that favour LD abundance.

To investigate, we performed transmission electron microscopy and observed extensive contacts between LDs and mitochondria in OA-supplemented cells **(Fig. 4A)**. The association between LDs and mitochondria also included ER structures as a frequent occurrence **(Fig. 4A)**. Quantitative analysis revealed that mitochondria were positioned significantly closer to the nearest LD in oleate-supplemented cells compared to control cells suggesting closer coupling of the two compartments **(Fig. 4B).** We next evaluated the frequency of LD-mitochondria contact sites and observed a marked increase both in the number of contact sites per cell as well as the length of the LD-mitochondria contact interface in LD-rich cells **(Fig. 4C, D)**. These observations are consistent with increased PDMs in oleate-treated cells. In agreement, a significantly higher proportion of mitochondria were detected within 30 nm of an LD in oleate-supplemented cells suggesting elevated PDM presence **(Fig. 4E)**. Notably, these changes in PDM occurrence were not due to differences in either the mitochondrial number or mitochondrial area per cell between control and LD-rich macrophages **(Fig. 4F, G)**. On the contrary, there was a modest but statistically significant decrease in the number of mitochondria under LD-rich conditions **(Fig. 4F)**. Consistent with the above observations, OA-supplemented cells exhibited a significant increase in LD size relative to control macrophages **(Fig. 4H)** suggesting that LD size may contribute to LD-mitochondria contact formation. In agreement, LD accumulation achieved through ATGL inhibition similarly enhanced LD size **(Fig. S6B)**. Conversely, exposure to palmitic acid and arachidonic acid had no significant effect on the size of LDs **(Fig. S4E)**. These data propose that LD size is coupled to peri-droplet mitochondria formation.

**Figure 4.**
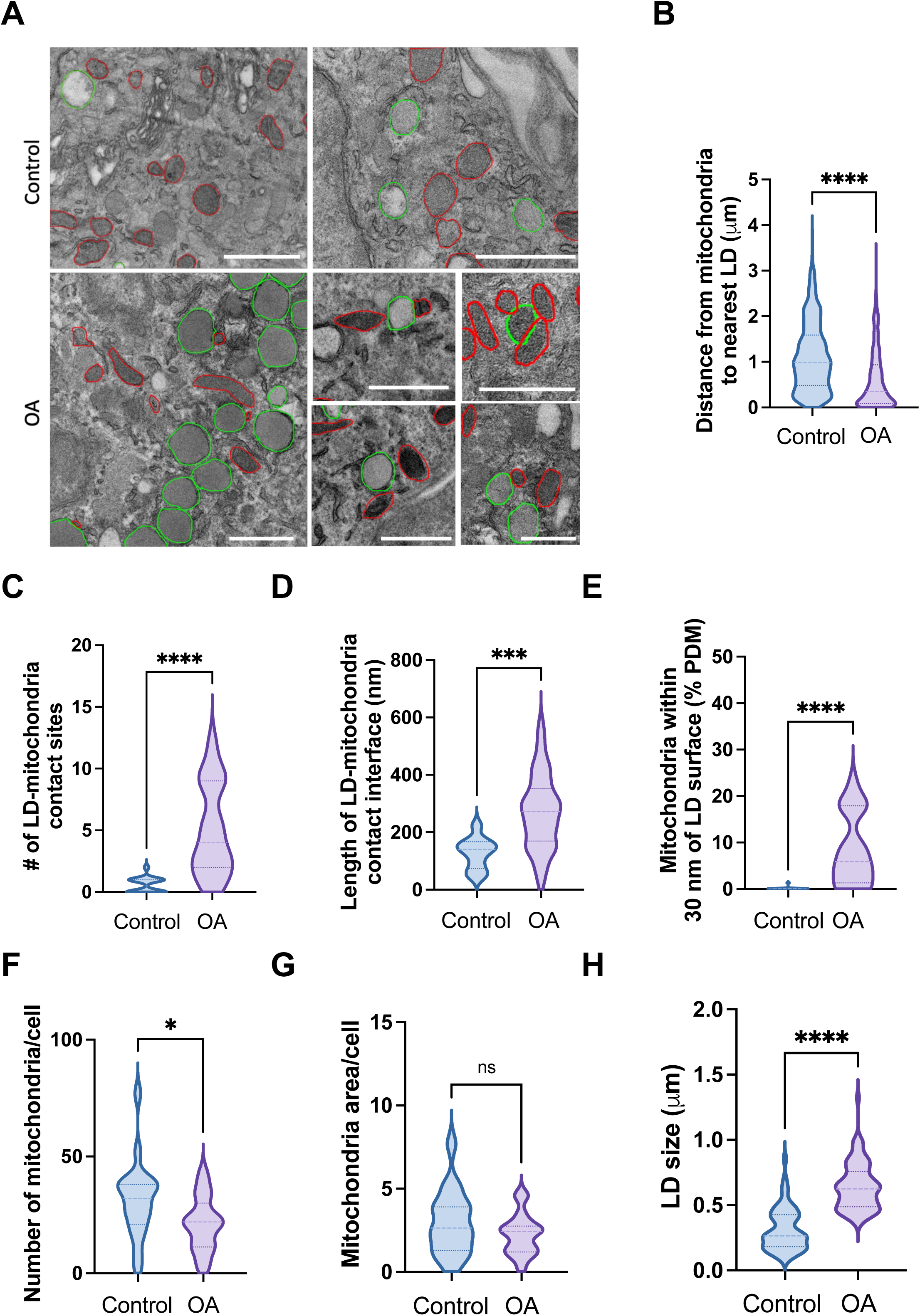
LD accumulation promotes increased peri-droplet mitochondria. **(A)** Transmission electron micrographs of BMDMs treated with LPS (500 ng/ml; 4 h) in the presence or absence of OA (90 µM; 24 h). LDs are outlined in green and mitochondria in red. **(B)** Quantitative analysis of mitochondria distance to nearest LD. **(C)** Quantitative analysis showing the number of LD-mitochondria contact sites, and **(D)** length of contact sites from cells treated as above. **(E)** Quantitative analysis by ImageJ of the percentage of mitochondria within 30 nm proximity to LD surface from analysis in (C). **(F)** Quantitative analysis showing the number of mitochondria, and **(G)** mitochondria area per cell in above treated cells. **(H)** Quantitative analysis by ImageJ of LD size in samples treated as above. Data shown are mean ± SEM, and experiments shown are representative of at least three independent experiments. Scale bars, 1 µM, *, p<0.05; ***, p < 0.001; **** P< 0.0001, by Student’s t test.

The two populations of mitochondria have also been shown to function differently and differ in structural aspects with PDMs more filamentous than cytoplasmic mitochondria. To investigate this, we labelled OA-supplemented macrophages with BODIPY (LDs) and MitoTracker (mitochondria) **(Fig. S7A)**. As expected, and in line with our earlier data, the fraction of mitochondria overlapping with LDs was significantly higher in the presence of expanded LDs **(Fig. S7B)**. Moreover, we observed increased mitochondrial branch length in cells supplemented with OA compared to control macrophages **(Fig. S7C)**. Altogether, these findings confirm that peri-droplet mitochondria are established during LD expansion, which may facilitate inflammasome activation.

### LD abundance drives mitochondrial remodelling and elevated oxidative metabolism

In response to increased substrate availability, cells undergo metabolic remodelling characterised by mitochondrial adaptation. To evaluate these adaptations in LD-rich cells, we labelled cells with membrane-potential sensitive dye MitoTracker Deep Red. Flow cytometry revealed that LD-rich cells exhibited an increase in membrane potential compared to untreated control macrophages **(Fig 5A, B)**. Notably, these changes were independent of mitochondrial ROS as MitoSox staining revealed comparable levels of superoxide production between control and LD-rich cells **(Fig. S8)**.

**Figure 5.**
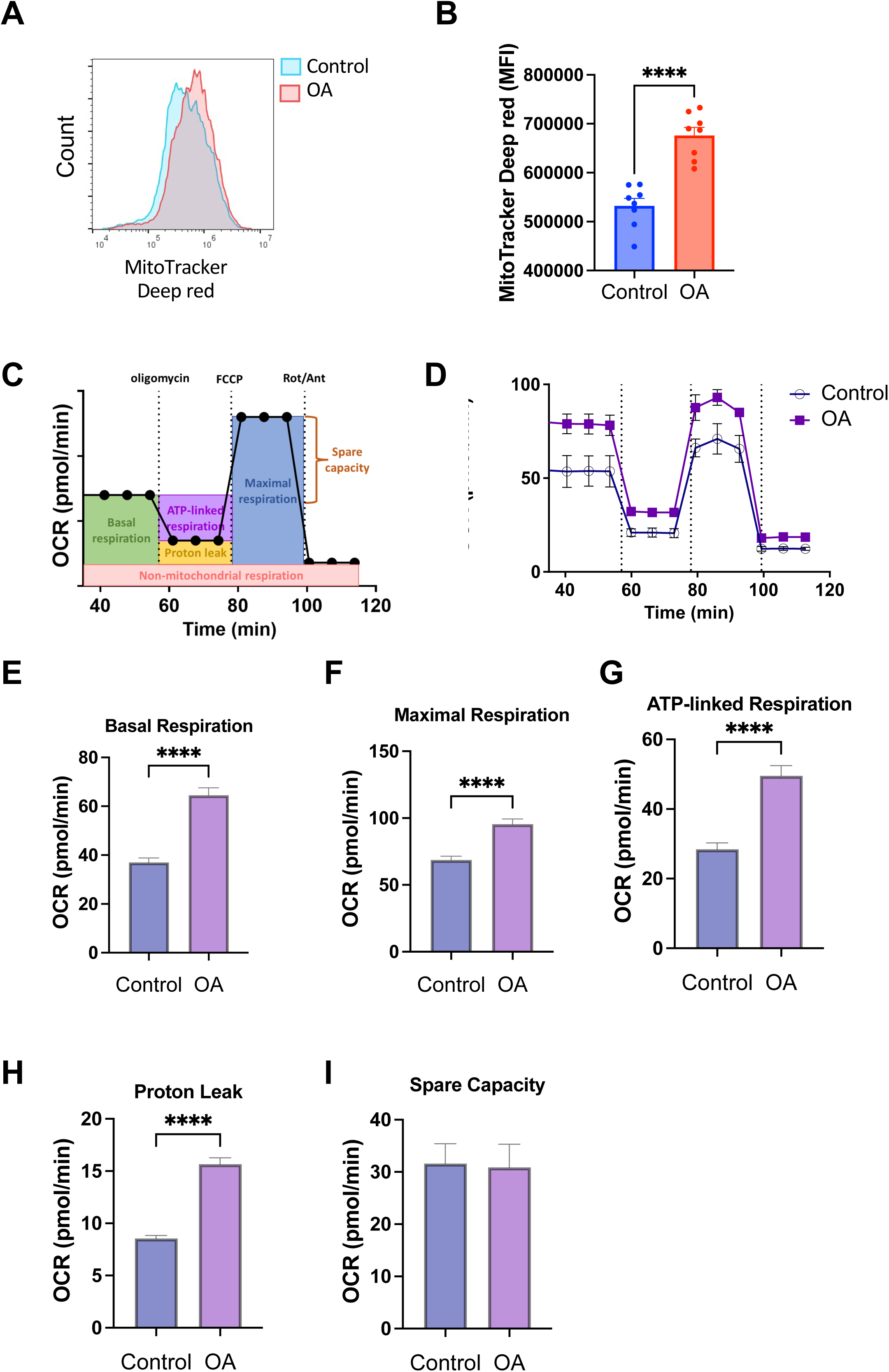
LD abundance drives mitochondrial remodelling and elevated oxidative metabolism. Immortalised BMDMs were treated with LPS (500ng/ml; 4 h) in the presence or absence of OA (30 µM; 48 h). **(A)** Representative histogram showing fluorescence intensity of MitoTracker Deep Red **(B)** Quantification of MitoTracker Deep Red MFI in the above treated cells. **(C)** Schematic representation of the Seahorse Mito Stress test showing the sequential **(D)** Representative traces from Seahorse Mito Stress test from above treated cells which were sequentially injected with oligomycin, FCCP, and Antimycin/Rotenone to assess mitochondrial oxygen consumption rate (OCR). **(E-G)** Quantification of basal, maximal, and ATP-linked respiration. **(H-I)** Quantification of proton leak and spare capacity. Data shown are mean ± SEM, and experiments shown are representative of at least three independent experiments. **** P< 0.0001, by Student’s t test.

Analysis of bioenergetics can provide important insights into how metabolic resources are utilised when cells adapt to available substrates. To next evaluate how LD accumulation influences cellular respiration, we utilised the Seahorse XF analyzer to measure the oxygen consumption rate (OCR) indicative of mitochondrial activity in real-time. Prior to the assay, we confirmed that equal cells were loaded in different wells. Compared to controls, real-time OCR profiling revealed clear differences in the respiratory behaviour of LD-rich macrophages. Cells containing LDs displayed significantly higher basal and maximal OCR, as well as increased ATP-linked respiration in an assay where pharmacological modulators of the electron transport chain were sequentially injected **(Fig. 5C-G)**. These results indicate increased oxidative phosphorylation (OXPHOS) capacity and elevated ATP-generating potential in LD-rich cells. Concurrently, these cells exhibited an increase in proton leak suggesting a mild uncoupling of the electron transport chain **(Fig. 5H)**. However, cells with LDs appeared to work closer to their maximal bioenergetic output and the spare capacity remained unchanged compared to control cells **(Fig 5I)**. Intriguingly, exogenous palmitic acid did not reveal major changes in mitochondrial bioenergetics while only a modest change was observed with arachidonic acid **(Fig. S9).** Together, these data reveal that LD expansion induces mitochondrial rewiring characterized by elevated mitochondrial activity and oxidative phosphorylation.

### Glutamine metabolism drives inflammasome activation in LD-rich macrophages

The ability of mitochondria to flexibly oxidise different substrates is central to maintaining immune and metabolic homeostasis. To further define substrate utilisation by LD-rich cells, we performed Seahorse substrate oxidation assays where we utilised selective inhibitors to block the mitochondrial utilisation of pyruvate, glutamine, or fatty acids. Remarkably, LD-rich cells demonstrated increased dependence on both pyruvate and glutamine oxidation **(Fig. 6A, B)**. By contrast, FAO reliance remained unchanged between control and LD-rich cells **(Fig. 6C)**. Intriguingly, supplementation with either palmitic acid or arachidonic acid resulted in no deviation from the use of pyruvate **(Fig. 6D)**. However, a decrease in fatty acid utilisation was observed in arachidonic acid supplemented cells **(Fig. 6E)**. These data suggest a shift in metabolic rewiring towards the use of alternative substrates, suggesting that LD accumulation drives mitochondrial bioenergetic flexibility.

**Figure 6.**
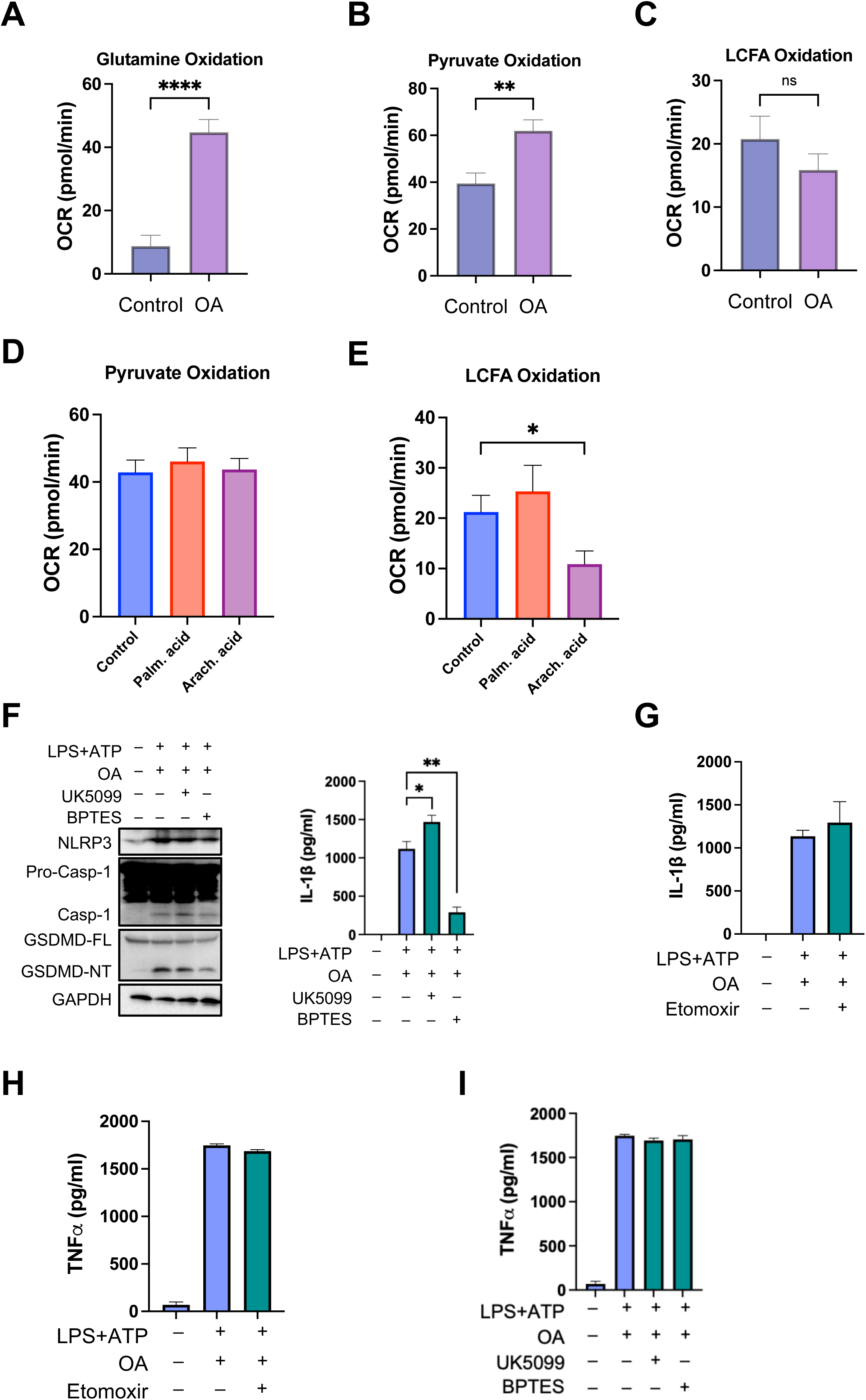
Glutamine metabolism drives inflammasome activation in LD-rich macrophages. Quantification of oxygen consumption rate (OCR) by control and OA-supplemented (30 µM; 48 h) iBMDMs using the Seahorse XF analyser to evaluate mitochondrial substrate oxidation following inhibition of the utilisation of individual substrates by mitochondria. **(A-C)** Plots depict OCR changes following inhibitor addition for glutamine, pyruvate, and long-chain fatty acids (LCFA) oxidation prior to FCCP uncoupling. **(D-E)** Plots depict OCR changes following inhibitor addition for pyruvate and long-chain fatty acids (LCFA) oxidation prior to FCCP uncoupling in palmitic and arachidonic acid supplemented cells. **(F, I)** Immunoblot showing expression of indicated proteins (*left panel*), IL-1β release (*right* panel), or TNF-α release **(I)** by OA-exposed (30 µM; 48 hrs) iBMDMs folowing LPS and ATP treatment in the presence or absence of inhibitors for either mitochondrial pyruvate carrier (UK5099, 30 µM; 24 h) or glutaminase (BPTES, 15 µM; 24 h). **(G, H)** IL-1β and TNF-α release by oleate-loaded cells in the presence or absence of inhibitor for carnitine palmitoyltransferase 1 (etomoxir, 24 h). Data shown are mean ± SEM, and experiments shown are representative of at least three independent experiments. **P< 0.01, **** P< 0.0001, by Student’s t test.

As the substrate oxidation assay above identified pyruvate and glutamine as major substrates in LD-enriched cells, we next examined whether these pathways directly influence inflammasome activation. To test this, cells were exposed to pharmacological inhibitors targeting either pyruvate or glutamine oxidation. Pyruvate oxidation relies on mitochondrial pyruvate carrier, which facilitates pyruvate import into mitochondria. Inhibition of mitochondrial pyruvate carrier by UK5099 resulted in a modest yet statistically significant increase in caspase-1 and GSDMD cleavage, and IL-1β secretion **(Fig. 6F)**. This may suggest that when mitochondrial pyruvate utilisation is restricted, alternative compensatory mechanisms may elevate inflammasome activation.

We next tested the role of glutamine metabolism in NLRP3 inflammasome activation under control and LD-rich conditions. Glutamine is sequentially converted to glutamate and α-ketoglutarate, which fuels the tricarboxylic acid cycle through glutaminolysis. The latter step is catalysed by the enzyme glutaminase (GLS1). Inhibition of glutaminolysis using the GLS1 inhibitor BPTES led to a marked reduction in the expression of cleaved forms of caspase-1 and GSDMD, and secreted IL-1β levels **(Fig. 6F)**. Additionally, and in agreement with the substrate oxidation data above, inhibition of FAO with etomoxir had no significant effect on either IL-1β or TNF-α secretion **(Fig. 6G, H)**. Notably, neither the inhibition of pyruvate oxidation nor glutaminolysis altered TNF-α release. These data confirmed that changes in IL-1β production are independent of inflammasome priming **(Fig. 6F)**. Importantly, these data demonstrate that glutaminolysis provides a specific bioenergetic input which is critical for inflammasome activation. Collectively, our study identifies glutamine metabolism as a critical metabolic driver of NLRP3 inflammasome activation in LD-loaded macrophages.

## Discussion

Despite emerging evidence linking lipid metabolism to NLRP3 activation, the roles of LDs in inflammasome activation have not been explored. Whether LD accumulation in macrophages primarily represents a metabolic adaptation to accommodate excess lipids or whether it also constitute a metabolic signal that actively regulates the NLRP3 inflammasome is not yet clear. Here, we identify LD abundance as a metabolic cue that potentiates NLRP3 inflammasome activation. We demonstrate that LD expansion, either achieved through exogenous oleic acid supplementation or the inhibition of lipolysis, strongly activates caspase-1 cleavage and IL-1β secretion. These observations causally link excess lipid storage to innate immune activation.

LD formation involves a balance between triglyceride synthesis and breakdown by lipolysis. Previous studies have demonstrated inconsistent results when examining the impact of these pathways on LD dynamics. Inhibition of either triglyceride synthesis or lipolysis both demonstrated blunted IL-6 secretion by macrophages^19,20^. Beyond lipid composition, LD identity is defined in part by the association of maturing LDs with different members of the perilipin protein family. For instance, perilipin 2, which is broadly expressed across cell types, is enriched in pro-inflammatory LDs associated with atherosclerotic lesions^50^. While perilipin 2 was not examined in our study, these observations suggest that the inflammatory nature of LDs can be governed by their molecular composition. The activation of NLRP3 inflammasome in atherosclerotic plaques has previously been linked to cholesterol crystals^15^. Our findings raise the possibility that LD accumulation within foamy macrophages may provide an additional trigger for inflammasome activation contributing to plaque progression.

A key finding in our study is the emergence of PDMs with elevated membrane potential and enhanced ATP-linked respiration in LD-rich macrophages. This physical coupling may facilitate the formation of metabolic microdomains that sustain bioenergetic output under LD-rich conditions. For example, we find that LD expansion unexpectedly redirects mitochondrial bioenergetic flux away from FAO. Instead, oleate-loaded cells utilised pyruvate and glutamine-derived carbon sources, suggesting a selective biochemical rewiring under LD-rich conditions. These metabolic changes may preserve mitochondria function while sustaining increased inflammatory function.

Surprisingly, our study demonstrates that glutaminolysis is a key metabolic pathway potentiating inflammasome activation. Dependent on GLS1, glutaminolysis results in the conversion of glutamine to glutamate and subsequently to α-ketoglutarate (α-KG)^51^. This metabolic intermediate eventually gets transferred to the TCA cycle to produce reducing equivalents (NADH and FADH_2_) and supports the generation of metabolites such as acetyl-CoA that provides both an energy source and carbons for various biosynthetic processes. We show that the disruption of glutaminolysis attenuates NLRP3 activation, highlighting the central role of glutamine metabolism in inflammasome activation. Glutaminolysis not only contributes to anaplerosis and ATP generation, but it also influences signalling intermediates such as α-ketoglutarate that regulate mitochondrial respiration and redox homeostasis. In agreement with the pro-inflammatory role reported in this study, glutamine metabolism was previously shown to calibrate mitochondrial ROS generation in neuroinflammation^52^. Moreover, GLS1 was upregulated in γδ T cells derived from patients with psoriasis, where it was shown to support Th17 and IL-17A-producing γδ T cell differentiation^53^. Accordingly, blocking GLS1 activity resolved Th17 and γδ T17 differentiation and epidermal hyperplasia in psoriasis-like mouse models^53^. Together, these studies suggest a broader role for glutaminolysis in inflammatory responses across diverse immune compartments.

Curiously, another study demonstrated an accelerated NLRP3 inflammasome activation when glutamine was deprived in the cell culture media^54^. This was due to reduced production of itaconate which suppresses NLRP3 inflammasome and may reflect context-specific metabolic conditions. Nonetheless, our study emphasises that glutaminolysis sustains mitochondrial bioenergetics efficiently aiding inflammasome activation in LD-rich cells.

Overall, our study has identified a previously unrecognised metabolic-inflammatory axis in which LD expansion and PDM formation regulate cellular metabolism through glutaminolysis to potentiate NLRP3 inflammasome activation. Targeting this axis may offer therapeutic opportunities to mitigate aberrant immune activation in metabolic disease and other chronic inflammatory conditions.

## Methods

### Ethics Statement

All animal procedures were carried out in accordance with the Animals (Scientific Procedures) Act 1986 and were approved by the Imperial College Animal Welfare and Ethical Review Body (AWERB) and the UK Home Office.

### Analysis of published transcriptomics data

Bulk RNA-seq data from wild-type C57BL/6 bone marrow-derived macrophages (BMDMs) were selected from GEO (GSE286554 [1], GSE224239 [2]). Experimental conditions were consistent with those described above: BMDMs were either primed with LPS and stimulated with ATP or treated with LPS alone, alongside untreated controls (n = 3 per group). Sequencing reads were processed through a reproducible pipeline comprising quality control, adapter trimming, alignment to Mus musculus GRCm39 reference genome, and gene-level quantification. Differential expression was performed with DESeq2 using Benjamini-Hochberg correction, and significant genes were defined as padj < 0.05 with |log_2_FC| > 1. Functional enrichment focused on lipid metabolic pathways, assessed with g:Profiler and summarized with *rrvgo* to reduce redundancy. Full details are provided in the **supplementary material (Supplementary files 1 to 7)**, and all analysis scripts are publicly available (https://github.com/ss-lab-cancerunit/Lipid_droplets).

### Cell culture

Mouse Bone marrow cells were isolated from 8–12-week-old mice femurs and tibia as previously described^55^. In brief, cells were allowed to differentiate into macrophages in DMEM containing 10% heat-inactivated FBS, 5% penicillin/streptomycin, 1% HEPES, 1% NEAA, and 30% L929 condition DMEM media. Cells were allowed to differentiate into bone-marrow derived macrophages (BMDMs) for 5-6 days at 37°C. immortalised BMDMs were cultured in DMEM containing 10% heat-inactivated FBS and 5% penicillin/streptomycin. Cells were grown in 10cm^2^ dishes and split every 2-3 days at approximately 90% confluency. THP-1 monocytes were cultured in RPMI containing 0% heat-inactivated FBS, 5% penicillin/streptomycin, 1% HEPES, 0.1% β-mercaptoethanol, and 2.5g/l extra D-glucose. THP-1 cells were grown in either T25 cm^2^ or T75 cm^2^ culture flasks and maintained at a density between 2×10^5^-1×10^6^ cells/ml and passaged every 3-4 days. THP-1 monocytes were differentiated into macrophages for 24 hrs with 20 nM phorbol-12-myristate-13-acetate (PMA) prior to cell stimulations. ATGL^−/−^ THP-1 monocytes were obtained from Synthego.

### Cell stimulations

Primary BMDMs, iBMDMs, and THP-1 monocytes were seeded in 24- or 12-well plates at a density of 0.5×10^6^ or 1×10^6^ cells, respectively. Experiments with OA (O3008; sigma-Aldrich) were optimised in different cell types to obtain sufficient LD accumulation. Primary BMDMs exposed to OA displayed sufficient LD accumulation at 24 h, while in iBMDMs sufficient LD accumulation was observed at 48 h. Palmitate and arachidonate were independently conjugated to fatty acid-free BSA prior to addition to cells. Where indicated, cells were exposed to palmitic acid or arachidonic acid for 24 h or 48 h before inflammasome activation in primary BMDMs or iBMDMs, respectively. Where used, cells were exposed to ATGListatin (SML1075; Sigma) 24 h prior to inflammasome activation at indicated concentrations.

### Inflammasome activation assays

For NLRP3 inflammasome activation, cells were primed with 500ng/ml LPS (tlrl-pbslps; Invivogen) for 3.5 h followed by either 5mM ATP (A6419; Sigma-Aldrich for 30-45 min or 10μM nigericin (4312; Tocris) for 1 h to 1.5 h. Pyroptosis cell images were taken following the addition of ATP using a light microscope. Where used, cells were activated using either alum at 1.7mg/ml or imiquimod at 30μg/ml 2 h after priming for 5-6 h. For NLRC4 inflammasome activation, cells were infected with *Salmonella typhimurium* at an MOI of 2 for 4 h. For AIM2 inflammasome activation, cells were stimulated with LPS 2 h prior to being transfected with 1μg of poly(dA:dT) using Lipofectamine 2000 (Invitrogen) according to manufacturer’s recommendation. Following each treatment, cell supernatants were collected and immediately frozen for cytokine analysis by ELISA, and cell lysates were harvested in radioimmunoprecipitation assay lysis (RIPA) buffer for immunoblot analysis.

### *In vivo* peritonitis model

The *in vivo* peritonitis model of inflammation was conducted on C57BL/6 mice (8–12-week-old) obtained from Charles River, UK. Animals were intraperitoneally injected with ATGListatin (50 mg kg−1; B3021; ApexBio) prepared in sesame oil. The next morning, animals were injected again with a second dose of ATGListatin prior to being intraperitoneally injected with LPS (100 μg kg^−1^). After 4 h, animals were injected with either PBS or ATP for 15 min. Mice were euthanised by carbon dioxide inhalation and approximately 500 μL of blood was immediately collected by cardiac puncture. Serum samples were stored at −20°C until cytokine levels in the serum were measured by ELISA.

For caspase-1 activity in vivo, peritoneal lavage cells were harvested in an equal volume (2.0 mL) of DMEM media before resuspending them in FACS buffer (1% BSA and 0.1% NaN^3^ in PBS) and transferring them to a 96 v-well plate. Cells were then washed once in FACS buffer and stained with FAM-FLICA Caspase-1 (YVAD) for 1 h at 37°C to label active caspase-1. Afterwards, cells were washed twice prior to staining and incubation with an antibody mix containing anti-mouse Ly-6G/Ly-6C (GR-1) antibody (0.25 mg/ml; Biolegend, 108424) and anti-mouse CD11b antibody (0.25 mg/ml; Biolegend, 101212) for 30 min at 4°C. After the incubation, cells were washed twice and resuspended in FACS buffer. Flow cytometry was performed on a Beckman Coulter’s Cytoflex, and raw data was analysed using the Flow-Jo software.

### Flow cytometry

To determine mitochondria membrane potential, cells were stained with mitochondrial membrane-dependent stain, MitoTracker Deep Red (100 nM; Thermo Fisher; M22426 in complete DMEM for 20 min at 37°C. After incubation, cells were washed twice and resuspended in FACS buffer (1% BSA and 0.1% NaN_3_ in PBS) before being transferred to 96-well v-bottom plate. To measure mitochondrial ROS, cells were stained with MitoSOX (5 μM; Thermo Fisher; M36008) in complete DMEM for 20 min at 37°C. All flow cytometry readings were performed on 96 v-well plates using a Beckman Coulter’s CytoFlex, and raw data were analysed using Flow-Jo Software.

### Confocal microscopy and Image analysis

Primary and immortalised BMDMs and HEK293T cells were plated on coverslips in 24-well plates. Coverslips for HEK293T cells were pretreated with poly-lysine solution for 10 min to support efficient adherence. For visualisation of LDs, following treatment, cells were washed in prewarmed PBS and stained with BODIPY 493/503 (2μM; Thermo Fisher; D3922) in PBS for 20 min at 37°C followed by washing twice in PBS. For mitochondria staining, cells were additionally stained with MitoTracker Deep red (100 nM; Thermo Fisher; M22426) in complete DMEM for 20 min at 37°C followed by washing twice in PBS. After staining with indicated stains, cells were fixed in 4% paraformaldehyde (PFA) for 30 min at room temperature. To remove autofluorescence caused by PFA, cells were quenched by being incubated with PBS-50mM glycine solution for 10 min at room temperature. After quenching, coverslips were mounted on slides with a drop of mounting media containing DAPI overnight at room temperature to stain the nucleus. For propidium iodide (PI) staining, cells on coverslips were stained at 500nM for 5 min followed by mounting coverslips on slides with DAPI mount. The slides were imaged on a SP5 confocal microscopy (Leica Microsystems), or a fluorescent EVOS microscope for PI images. Images were processed and analysed on ImageJ. For mitochondria analyses, Iterative Deconvolve 3D, skeletonize (2D/3D), and MiNa Mitochondrial Network Analysis plugins were used.

### Electron microscopy

Treated cells were fixed in 2% PFA and 2% glutaraldehyde fixative in 0.1 M cacodylate buffer for 30 min at room temperature. After fixation, coverslips were washed in 0.1 M cacodylate buffer and post-fixed in osmium tetroxide for 1 h at 4°C in the dark. To enhance contrast, cells were incubated in 1% tannic acid in 0.05 M cacodylate buffer for 40 min at room temperature. Cells were then washed in 0.1 M cacodylate buffer and dehydrated in a descending series of graded ethanol and finally embedded in 1:1 of propylene oxide and epon resin mixture which was later replaced by neat epon. Cells were then mounted on Epon stubs and baked at 65°C overnight. Samples were trimmed and cut en face at 70 nm thickness with a diamond knife (DiATOME) in a Leica Ultracut UCT 6 ultramicrotome before examination on an FEI Tecnai G2-Spirit TEM. Images were acquired in a charge-coupled device camera (Eagle) and analysed in Fiji.

### Seahorse XF analysis

All oxygen consumption rate (OCR) and extracellular acidification rate (ECAR) were conducted using Agilent Seahorse XF^e^96 analyser (Seahorse Biosciences). In brief, cells were seeded at a density of 40,000 cells per well and allowed to adhere to a 96-well XF cell culture microplate (Seahorse Biosciences). For Mito stress test, sequential OCR and ECAR measurements were taken after the addition of oligomycin (1.5μM), FCCP (1μM), and rotenone (0.5μM) plus antimycin A (0.5μM) at indicated timepoints. The measurements were used to calculate basal respiration, maximal respiration, ATP-linked respiration, proton leak, and spare capacity according to manufacturer’s instructions. Basal ECAR measurements were plotted as approximate basal glycolysis. For substrate oxidation test, where indicated, UK5099 (15μm), BPTES (10μM), or Etomoxir (20μM) were injected in port A to calculate pyruvate, glutamine, or FAO, respectively, as per manufacturers recommendation. Control wells were injected with seahorse assay media instead.

### Immunoblotting and antibodies

For immunoblotting, samples were first boiled at 95°C before resolving them on 12% SDS-page gels and transferring them onto nitrocellulose membranes (GE Life Sciences). Membranes were blocked for 1 h at room temperature with 5% milk solution in TBS containing 0.05% Tween-20. Membranes were then incubated with primary antibodies at 4°C overnight. HRP-conjugated secondary antibodies were used for 1h at room temperature. Proteins were visualised using the Clarity ECL substrate (Bio-Rad) or the SuperSignal West Femto ECL substrate (ThermoFisher). Images were acquired on a BioRad imager and processed using Image Lab (Bio-Rad). The primary antibodies were used as follows: Mouse and human anti-caspase-1, mouse anti-NLRP3 and mouse anti-ASC were acquired from Adipogen, anti-GAPDH (Life Technologies), pro-IL-1b (Cell Signaling), cleaved IL-1b (Cell Signaling), ATGL (Cell Signalling), GSDMD (Abcam), and PLIN5 (Proteintech). Secondary antibodies were obtained from ThermoFisher Scientific and used at a dilution of 1:5,000.

### Enzyme-linked immunosorbent assay

Cell culture supernatants or serum were measured for mouse IL-1β (88-7013-88; Invitrogen), human IL-18 (RAB0543A; Sigma), and mouse TNFα (88-7324-88; Invitrogen) using ELISA kits according to the manufacturer’s instructions.

### Quantification & Statistical Analysis

GraphPad Prism 10.0 software was used for data analysis. Data are represented as mean ± SD or SEM (as indicated in the figure legends) and are representative of experiments done at least three times. Statistical significance was determined by unpaired Student’s t test or one-way ANOVA; p < 0.05 was considered statistically significant.

## Acknowledgements

We are grateful to Tristan Rodriguez for helpful comments on the manuscript. This work was in-part supported by a grant from The Medical Research Council, UK (MR/S00968X/1) to P.A. The research in the lab of P.A is funded by MRC grant (UKRI1408). The A.T. lab is supported by a project grant from The Medical Research Council (MR/X021467/1) and a Discovery Award from the Wellcome Trust (301619/Z/23/Z). F.W.K.T. is supported by the Ken and Mary Minton Chair of Renal Medicine (Grant number G30614).

The funders had no role in the design, conduct, or preparation of this manuscript.

## Author Contributions

N.A., A.O., and P.K.A. contributed to experimental design; N.A., E.P., Y.Z., G.A., A.O., and A.T. performed experiments; N.A., E.P., Y.Z., and P.K.A. analysed data; G.F., S.S., A.T., and F.W.K.T. participated in discussions, provided intellectual input, and suggested edits to the manuscript; P.K.A. designed the study, provided resources and overall supervision, and wrote the original draft. All authors approved the final version.

