## Supplementary material for "Lipid droplet accumulation drives glutamine-dependent NLRP3 inflammasome activation": Fig. S

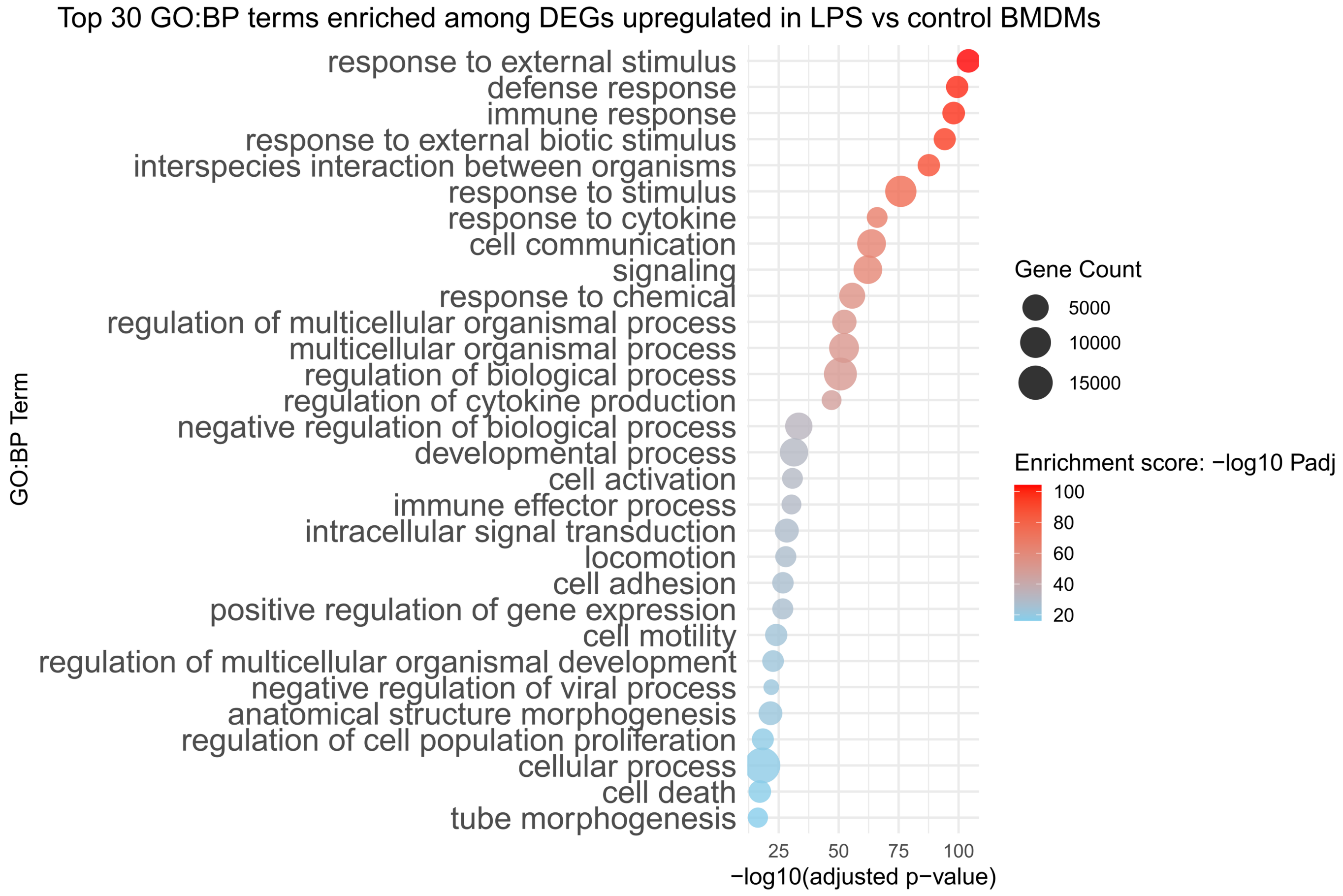


**Figure S1. NLRP3 inflammasome-activating stimuli remodel lipid metabolism and storage pathways, related to figure 1.**

Top 30 enriched Gene Ontology Biological Process (GO:BP) terms among upregulated DEGs in LPS treated versus control BMDMs, identified by g:Profiler and summarized with rrvgo. Enrichment score = -$\log_{10}$ (FDR-adjusted p value); gene count = number of DEGs within each GO:BP term. Full results are provided in Supplementary Materials.

**Figure S1**

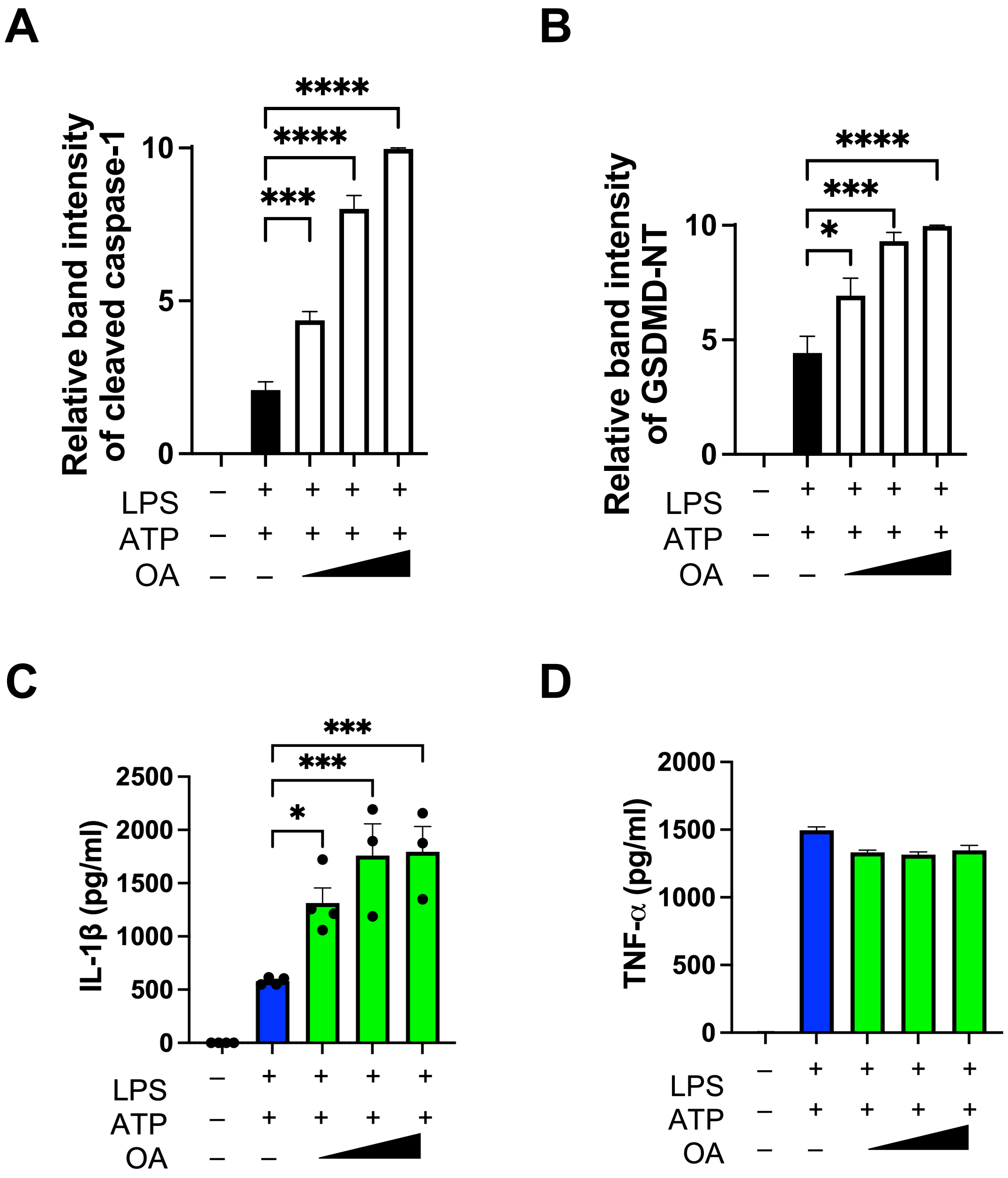


**Figure S2. LD accumulation results in increased NLRP3 inflammasome activation and pyroptosis, related to figure 2.**

Immortalised BMDMs stimulated with inflammasome priming signal LPS (500 ng/ml; 3.5 hours) were exposed to increasing concentrations of OA (30, 60, and 90 μM) for 48 h prior to exposure to ATP (5 mM; 30 min).
**(A, B)** Relative band intensity of cleaved caspase-1 and cleaved GSDMD as measured by ImageJ.
**(C, D)** IL-1β and TNF-α release measured by ELISA.
Data shown are mean ±SD. Experiments shown are representative of at least three independent experiments. *, $p<0.05$; ***, $p < 0.001$; ****, $p < 0.0001$, by one-way ANOVA, or Student's t test.

**Figure S2**

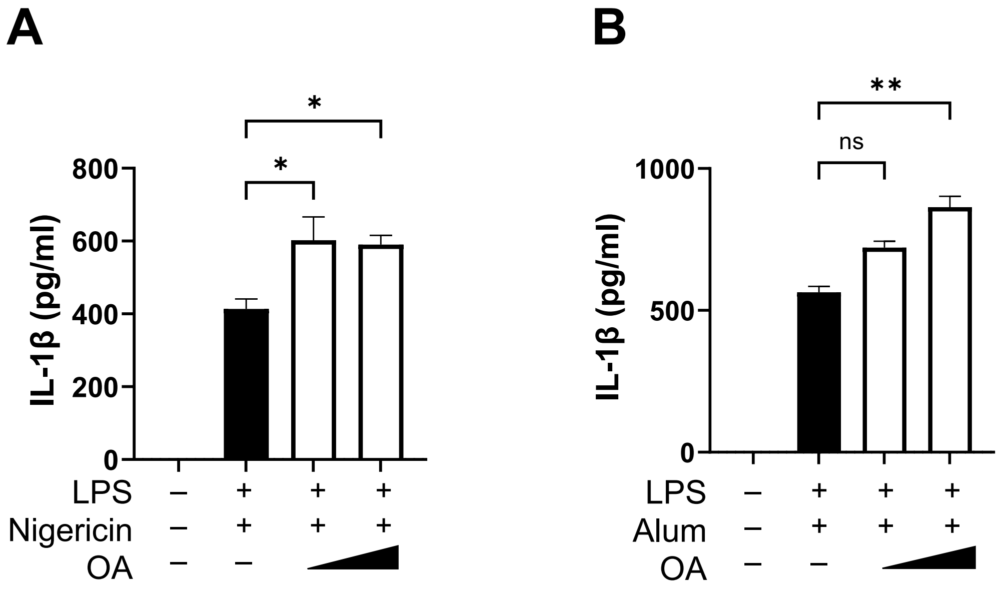


**Figure S3. LD accumulation results in increased NLRP3 inflammasome activation and pyroptosis, related to figure 2.**

**(A)** Immortalised BMDMs stimulated with inflammasome priming signal LPS (500 ng/ml; 3.5 h) were exposed to increasing concentrations of OA (30, 60, and 90 µM) for 48 h prior to exposure to Nigericin (10 µM; 1 h). IL-1β release measured by ELISA.
**(B)** Immortalised BMDMs stimulated with inflammasome priming signal LPS (500 ng/ml; 3.5 h) were exposed to increasing concentrations of OA (30, 60, and 90 µM) for 48 h prior to exposure to alum (1.7 mg/ml; 5 h). IL-1β release measured by ELISA.
Data shown are mean ±SD. Experiments shown are representative of at least three independent experiments. *, $p<0.05$; **, $p < 0.01$, by Student's t test.

### Figure S3

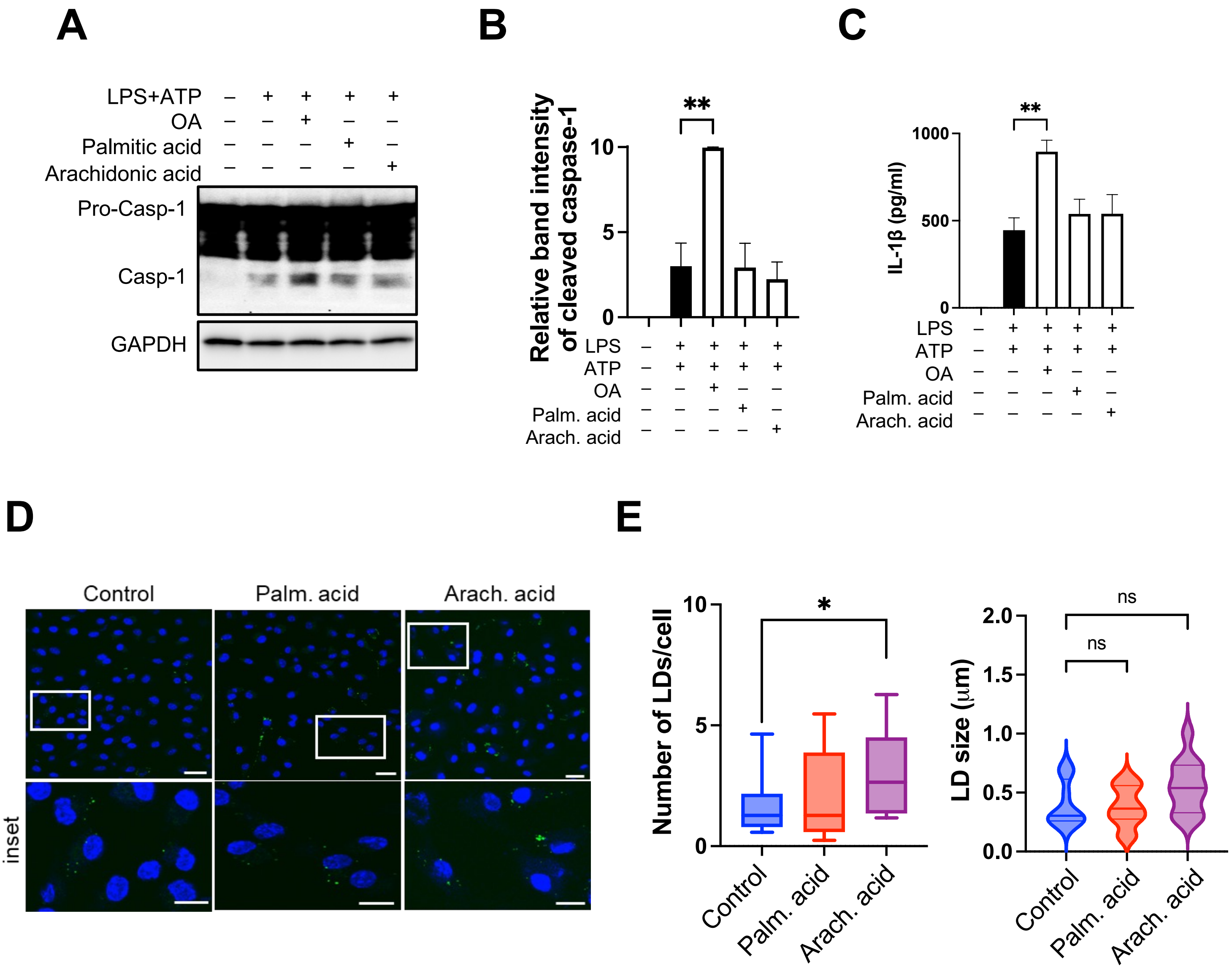


**Figure S4. LD accumulation results in increased NLRP3 inflammasome activation and pyroptosis, related to figure 2.**

**(A)** BMDMs stimulated with inflammasome priming signal LPS (500 ng/ml; 3.5 hours) were exposed to either OA (90 μM), palmitic acid (90 μM), or arachidonic acid (5 μM) for 24 h prior to exposure to ATP (5 mM; 30 min). Cell lysates were immunoblotted with the indicated antibodies.
**(B)** Relative band intensity of cleaved caspase-1 as measured by ImageJ.
**(C)** IL-1β release measured by ELISA from cells treated as above.
**(D)** Confocal microscopy images of BMDMs grown in the presence or absence of palmitic acid (90 μM), or arachidonic acid (5 μM) for 24 h followed by incubation with BODIPY 493/503 (2 μM for 20 min in PBS) at 37°C.
**(E)** Quantitative analysis of the number of lipid droplets and LD size in cells treated as above.
Data shown are mean ±SD. Experiments shown are representative of at least three independent experiments. *, $p<0.05$; **, $p < 0.01$ by one-way ANOVA, or Student's t test.

### Figure S4

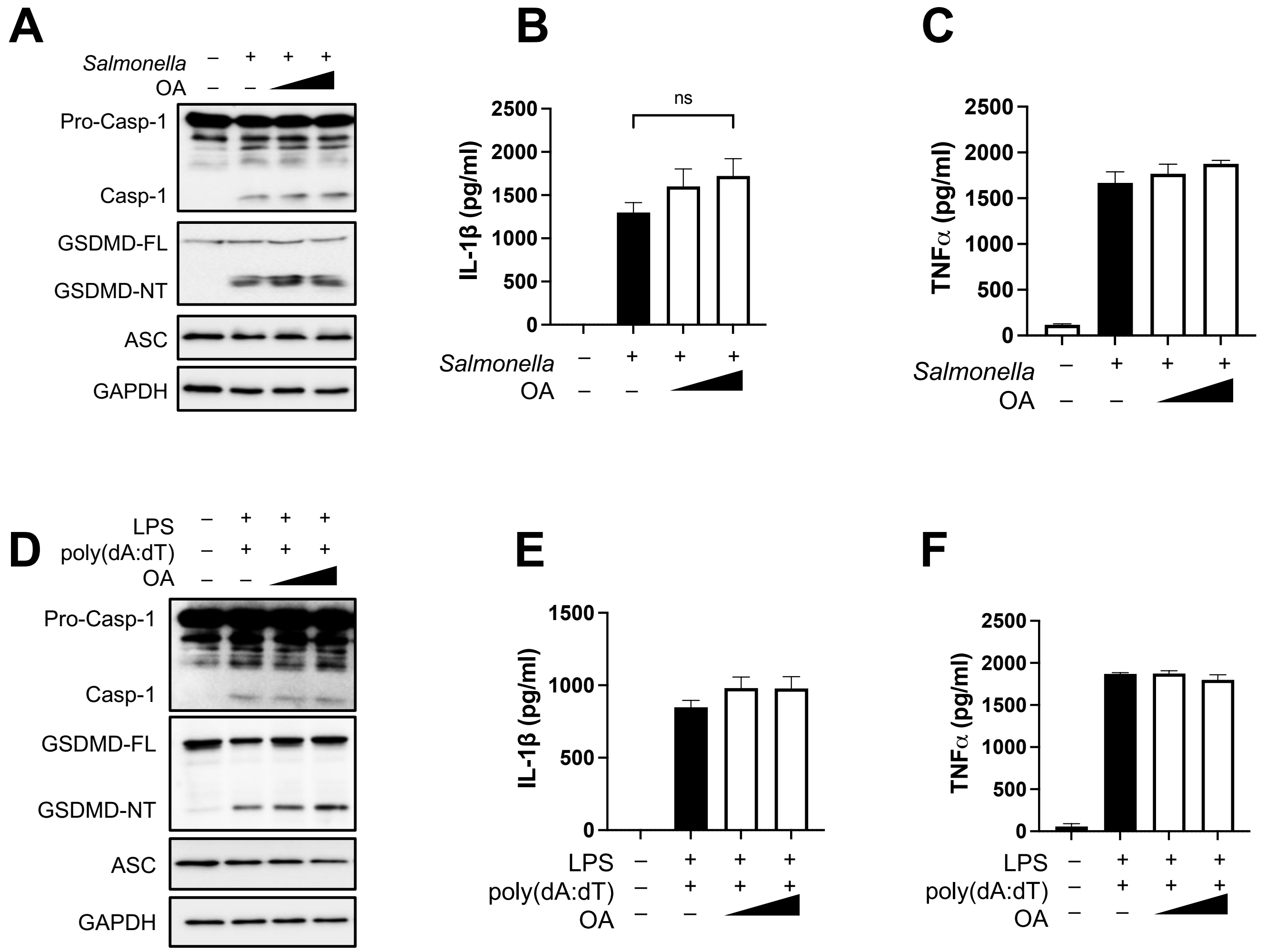


**Figure S5. LD accumulation does not affect the activation of NLRC4 and AIM2 inflammasomes.**

**(A)** Immortalised BMDMs were infected with *S. typhimurium* at an MOI of 2 for ~6 hours with or without the presence of increasing concentrations of OA (15, 30, and 45 μM; 48 h). Cell lysates were collected and used to visualise the proteins indicated.
**(B-C)** IL-1β and TNF-α release measured by ELISA from cells treated as above.
**(D)** Immortalised BMDMs stimulated with LPS (500 ng/ml; 2 h) were exposed to increasing concentrations of OA (15, 30, and 45 μM; 48 h) following transfection with the AIM2 agonist, poly (dA:dT) for 8 hrs. Cell lysates were collected and used to visualise the proteins indicated.
**(E-F)** IL-1β and TNF-α release measured by ELISA from cells treated as above.
Data shown are mean ±SD, and experiments shown are representative of at least three independent experiments.

### Figure S5

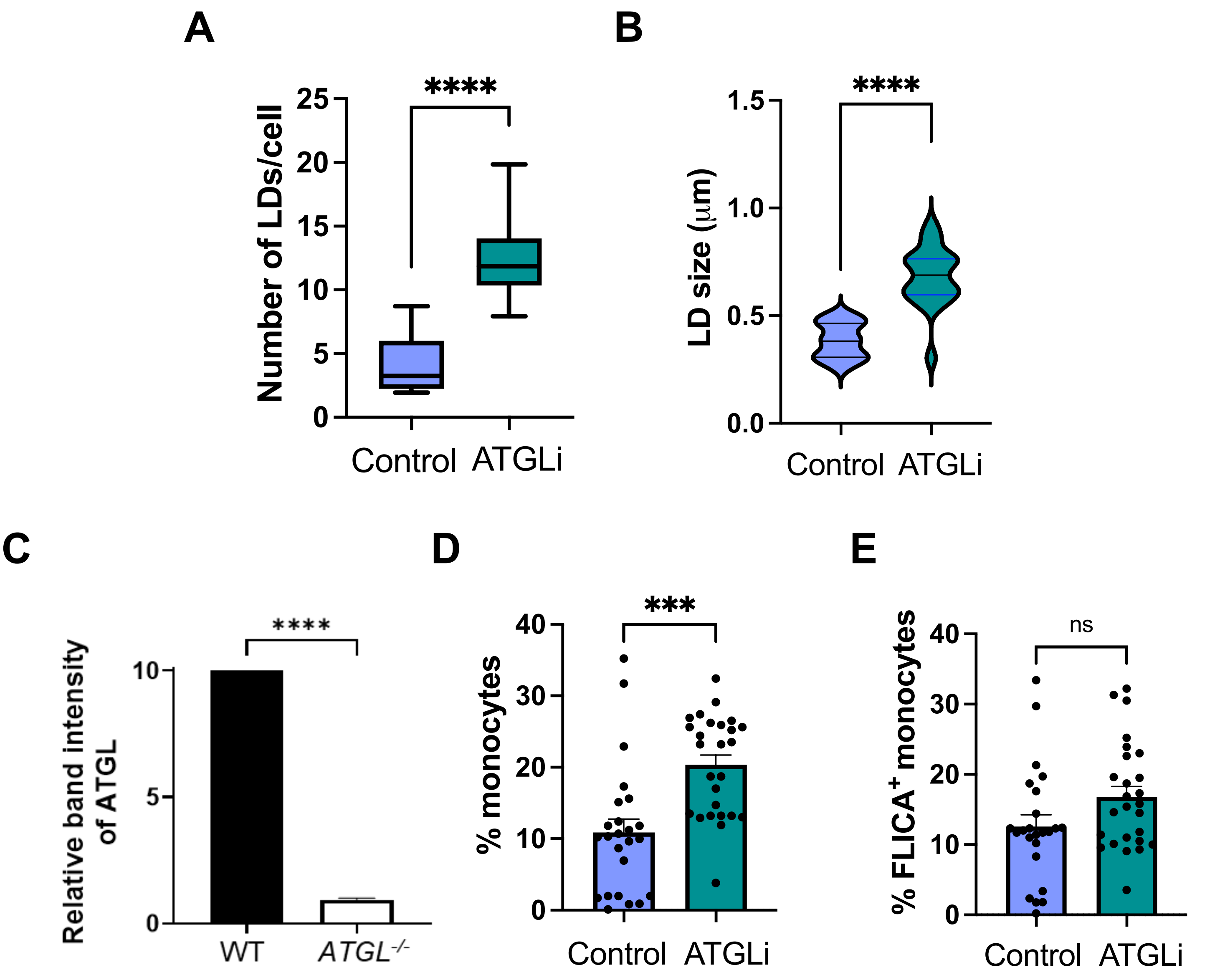


**Figure S6. Inhibition of lipolysis elevates NLRP3 inflammasome activation in vitro and in vivo, related to figure 3.**

**(A-B)** Confocal microscopy images of BMDMs grown in the presence or absence of ATGL inhibitor (ATGLi; 40 µM; 24 h) followed by incubation with BODIPY 493/503 (2 µM for 20 min in PBS) at 37°C. Quantitative analysis of the number of lipid droplets (LDs) per cell and LD size.

**(C)** WT and *ATGL*$^{-/-}$ THP-1 macrophages differentiated with PMA. Relative band intensity of ATGL as measured by ImageJ.

**(D-E)** C57BL/6 mice were administered intraperitoneally (i.p.) either with vehicle control (n=12) or ATGLi (50 mg kg−1; n=13) and fasted for 16 h. Next day, mice were again administered either vehicle control or ATGLi and LPS (100 µg kg−1). After 4 h, mice were i.p. injected with ATP for 15 min. Frequency of total CD11b+Gr1+ monocytes and caspase-1 active monocytes in peritoneal exudate cells.

Data shown are mean ±SD. Experiments shown are representative of at least three independent experiments. ***, $p < 0.001$; ****, $p < 0.0001$, by Student's t test.

**Figure S6**

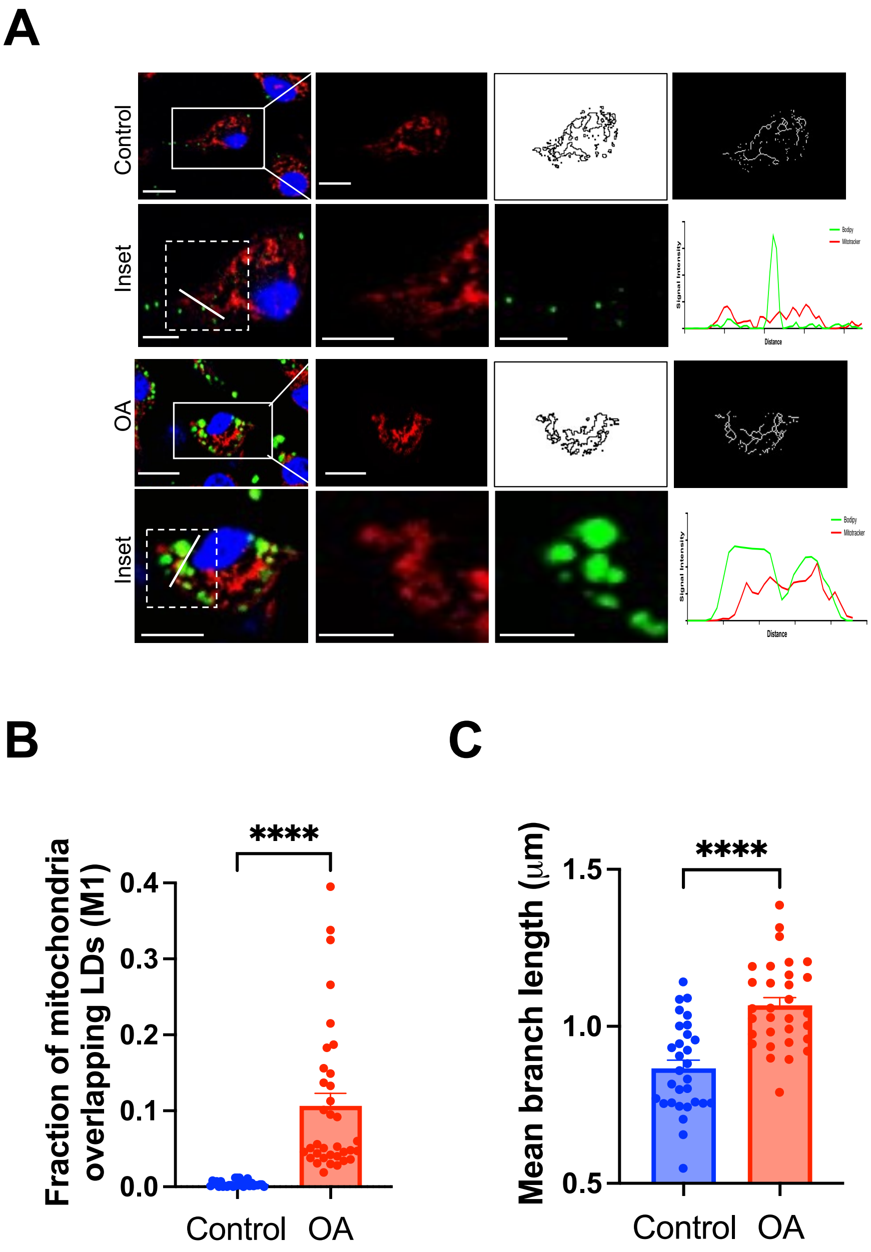


**Figure S7. LD accumulation promotes increased peri-droplet mitochondria, related to figure 4.**

**(A)** Confocal microscopy images of BMDMs grown in the presence or absence of OA (90 µM; 24 h) followed by incubation with BODIPY 493/503 (2 µM for 20 min in PBS) and MitoTracker Deep Red (100nM for 20 min in complete DMEM) at 37°C. Mitochondrial morphology analysis showing mitochondrial outline and skeleton performed by ImageJ. Profile of fluorescent intensity plot along the indicated bars in inset images.
**(B)** Quantitative plot representing fraction of MitoTracker overlapping BODIPY using Mander's coefficient (M1).
**(C)** Quantification of mean mitochondrial branch length as measured by ImageJ.
Data shown are mean ±SD. Experiments shown are representative of at least three independent experiments. ****, $p < 0.0001$, by Student's t test.

### Figure S7

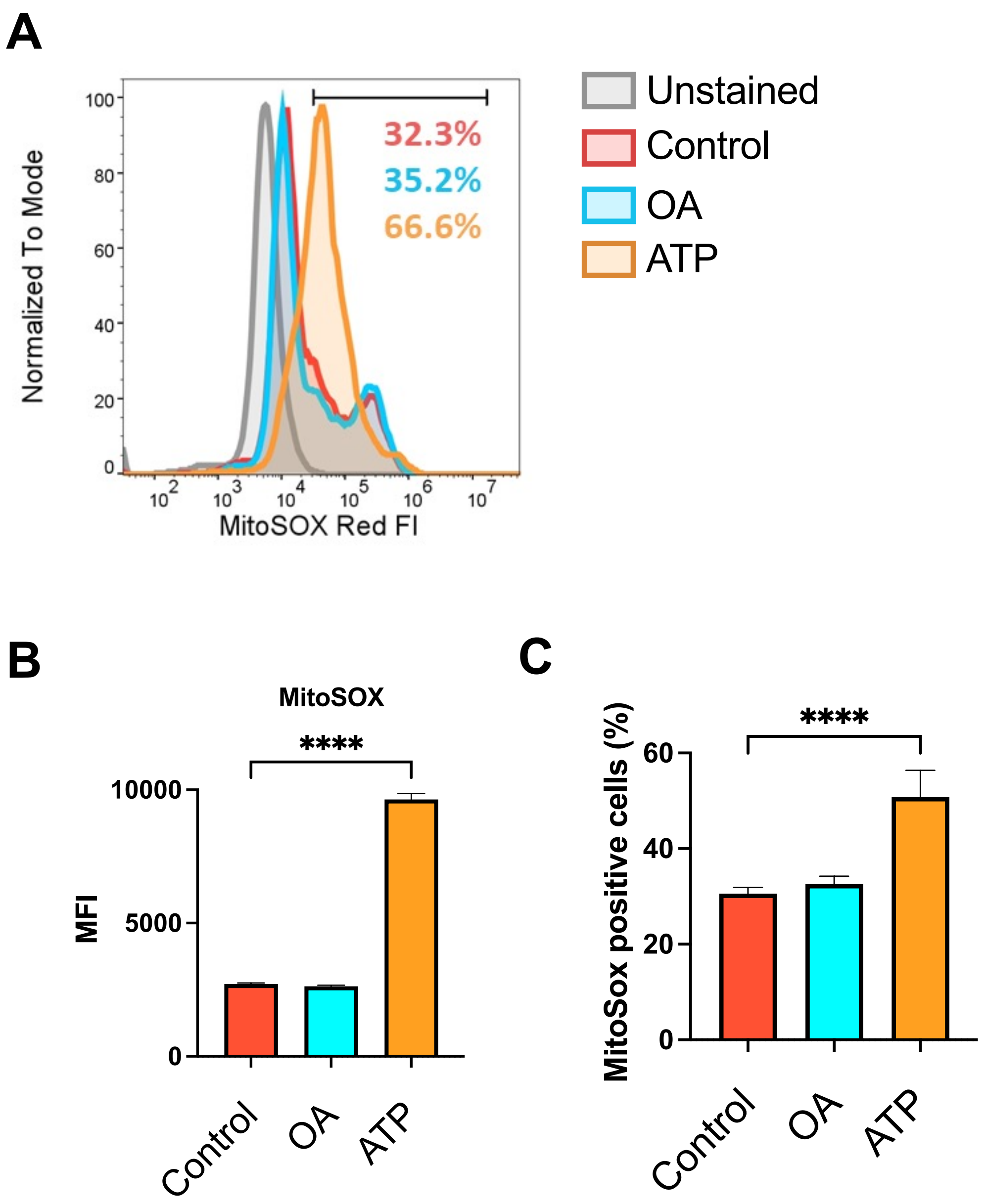


**Figure S8. LD accumulation promotes increased peri-droplet mitochondria, related to figure 4.**

BMDMs grown in the presence or absence of either OA (90 μM; 24 h) or ATP (5mM; 30 min) followed by incubation with MitoSOX (5 μM for 20 min in complete DMEM) at 37°C.
**(A)** Representative histogram showing fluorescence intensity of MitoSOX.
**(B-C)** Quantification of MitoSOX MFI and % MitoSOX positive cells in the above treated cells.
Data shown are mean ±SD. Experiments shown are representative of at least three independent experiments. ****, $p < 0.0001$, by Student's t test.

### Figure S8

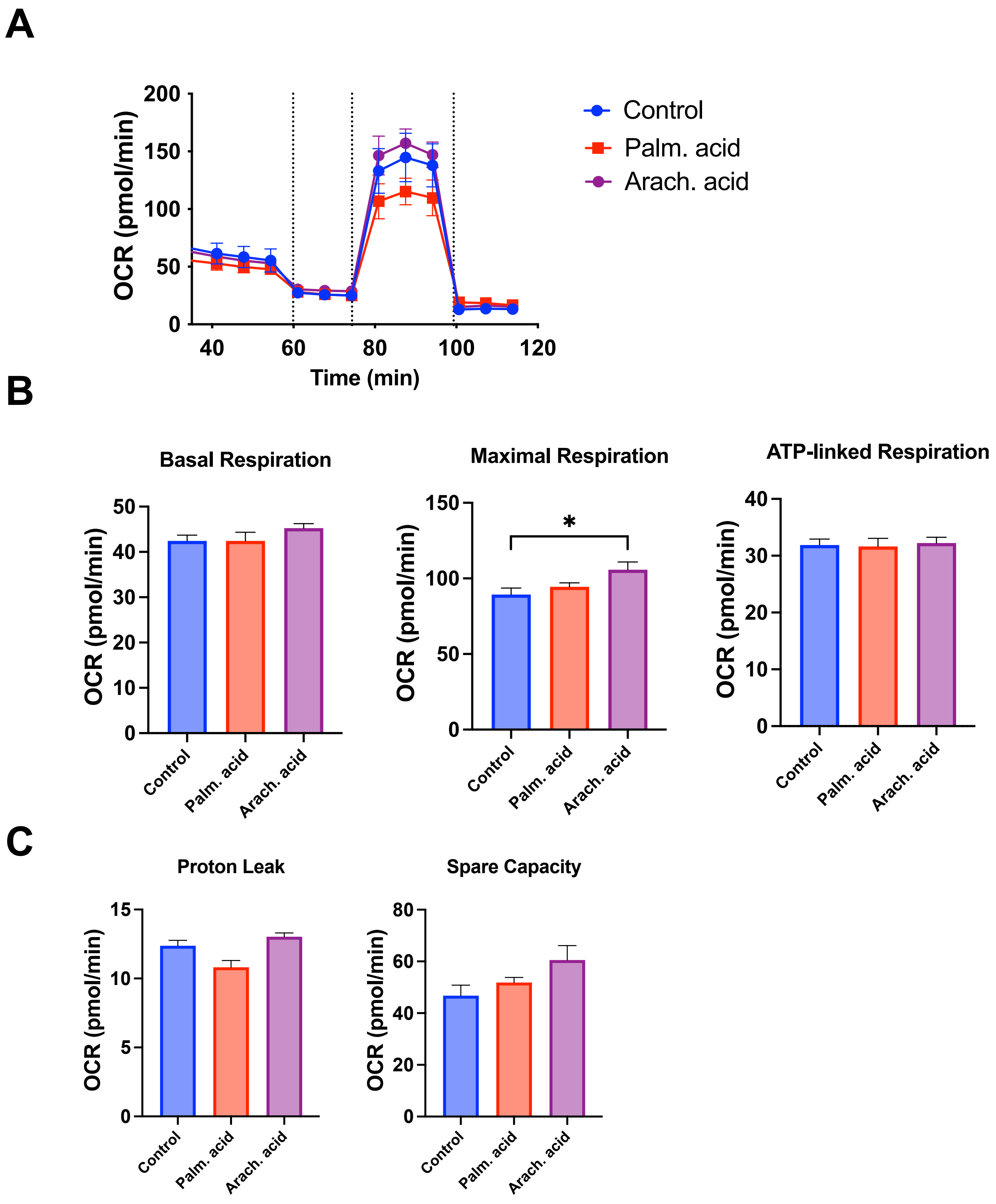


**Figure S9. Palmitic acid and arachidonic acid do not drive mitochondrial remodelling.**

**(A)** Immortalised BMDMs grown in the presence or absence of either palmitic acid (90 µM; 24 h) or arachidonic acid (5 µM; 24 h). Representative traces from Seahorse Mito Stress test from above treated cells which were sequentially injected with oligomycin, FCCP, and Antimycin/Rotenone to assess mitochondrial OCR.
**(B)** Quantification of basal, maximal, and ATP-linked respiration.
**(C)** Quantification of proton leak and spare capacity.
Data shown are mean ±SD. Experiments shown are representative of at least three independent experiments. * $p<0.05$ by Student's t test.

**Figure S9**
